# Separation estimation of two freely rotating dipole emitters near the quantum limit

**DOI:** 10.1101/2025.01.03.631147

**Authors:** Sheng Liu, Sujeet Pani, Sajjad A. Khan, Francisco E. Becerra, Keith A. Lidke

## Abstract

According to Rayleigh’s criterion, two incoherent emitters with a separation below the diffraction limit are not resolvable with a conventional fluorescence microscope. One method of Super-Resolution Microscopy (SRM) circumvents the diffraction-limited resolution by precisely estimating the position of spatiotemporally independent emitters. However, these methods of SRM techniques are not optimal for estimating the separation of two simultaneously excited emitters. Recently, a number of detection methods based on modal imaging have been developed to achieve the quantum Cramér-Rao lower bound (QCRB) to estimate the separations between two nearby emitters. The QCRB determines the minimum achievable precision for all possible detection methods. Current modal imaging techniques assume a scalar field generated from a point source, such as a distant source from an optical fiber or a pinhole. However, for fluorescently labeled samples, point emitters are single fluorophores that are modeled as dipole emitters and, in practice, are often freely rotating. Dipole radiation must be described by vectorial theory, and the assumption of a scalar field no longer holds. Here, we present a method to numerically calculate the QCRB for measuring the separation of two dipole emitters, incorporating the vectorial theory. Furthermore, we propose a near-quantum optimal detection scheme based on one of the modal imaging techniques, super-localization by image inversion interferometry (SLIVER), for estimating the separation of two freely rotating dipoles. In the proposed method, we introduce a vortex wave plate before the SLIVER detection to separate the radial and azimuthal components of the dipole radiation. With numerical simulations, we demonstrated that our method achieves non-divergent precision at any separation between two dipole emitters. We investigated several practical effects relevant to experimental measurements in super-resolution microscopy, including numerical aperture, detection bandwidth, number of estimation parameters, background, and misalignment on separation estimation. Our proposed measurement provides a near quantum-limited detection scheme for measuring the separation of two freely-rotating dipole emitters, such as fluorescently tagged molecules, which are commonly used in super-resolution microscopy.

## 1. Introduction

Rayleigh’s criterion is commonly used to assess the resolution limit of an imaging system and defines the minimum separation at which two emitters are resolvable [1]. This resolution limit is termed the diffraction limit, which can be approximated as the width of the impulse response of an imaging system, the point spread function (PSF) [2]. Given the output response and the PSF model for a particular imaging system, the estimation precision of a set of unknown parameters is bounded by the classical Cramér-Rao lower Bound (CRB) [3, 4]. For example, the CRB is widely used in single-molecule localization microscopy (SMLM) to determine the localization precision of a point emitter [5–11]. The detection methods for most SMLM systems can be categorized as direct detection, where the electric field from a point source is directly detected without any decomposition. For direct detection of two emitters, the CRB for separation estimation diverges quickly toward infinity as the separation decreases much below the diffraction-limited resolution [12, 13].

While the CRB depends on the particular measurement of an unknown parameter, quantum information theory provides a bound for any possible measurement of the parameter allowed by quantum mechanics, the quantum CRB (QCRB) [12, 14–17]. This relationship can be expressed as

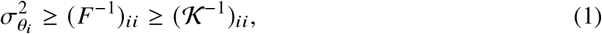

where *F* denotes the classical Fisher information (FI) matrix of a particular measurement on a set of parameters *θ. σ*^2^ denotes the variance of an estimator. 𝒦 denotes the quantum Fisher information (QFI) matrix of measuring the same set of parameters *θ* given a particular quantum state (e.g., a single-photon state, a squeezed state, an entangled photon state, etc.). Importantly, 𝒦 depends on the quantum state and is independent of any particular measurement strategy, i.e. optimized over all measurements. Finding a measurement strategy that saturates this bound is a fundamental goal in quantum metrology [18, 19].

In the case of fluorescence imaging, the fluorophore can be considered as a weak thermal source, whose quantum state can be approximated to an incoherent mixture of vacuum (zero-photon) and one-photon states [12]. As the vacuum state provides no information, all information for estimating a parameter is encoded in the one-photon state. For the case of two Gaussian emitters, the QFI for separation estimation is *N* /4*σ*^2^, with *N* being the expected photons from the two emitters and *σ* the width of the Gaussian [12]. This result shows that the QFI for separation estimation of two Gaussian emitters is independent of the emitters’ separation.

Furthermore, Tsang et al. proposed a measurement technique that achieves the QCRB for estimating the separation of two scalar point sources [12, 20, 21]. This measurement technique is based on modal imaging, which measures the projection of the input field onto a set of orthogonal spatial modes, such as Hermite-Gaussian modes. Measurement of the number of photons in each mode can be used in an estimator that achieves the QCRB. Several experimental realizations of modal imaging have been demonstrated. One implementation uses a spatial mode demultiplexing (SPADE) device (or a mode sorter) to decompose the input field onto a set of orthogonal spatial modes [22,23]. Some implementations project the input field onto fewer spatial modes. For instance, super-resolved position localization by inversion of coherence along an edge (SPLICE) projects the input field onto a single spatial mode, which is generated by introducing a *π* phase shift between the top and bottom halves of the input field [24]. Alternatively, by spatially inverting the input field in one arm of a Mach-Zehnder interferometer, super-localization by image inversion interferometry (SLIVER) decomposes the input field into its symmetric and antisymmetric components. SLIVER was first proposed by Nair et al. [25], and later experimentally demonstrated by Tang et al. [26].

However, all previous implementations of modal imaging were demonstrated by using a fiber-coupled laser as a point source. Such point sources can be described by scalar PSFs, such as a Gaussian PSF or an Airy pattern [2]. In contrast, in the field of fluorescence imaging, point sources are single fluorophores, which are dipole emitters [11, 27–35]. The PSF of a static dipole emitter is described by vectorial diffraction theory, and the PSF pattern varies with the dipole orientation. Furthermore, in many fluorescently labeled samples, the fluorophore can freely rotate in an aqueous medium. We can thus model the fluorophore as a freely rotating dipole, where the dipole rotation is much faster than the fluorescent lifetime, causing the resulting radiation to sample all possible dipole orientations [29]. For modeling the PSF of a freely rotating dipole, each emitted photon is considered as originating from a static dipole, and the response of the optical system is treated with vectorial diffraction theory. In summary, the quantum state of a dipole emitter consists of a mixture of fields from dipoles oriented in different directions. Previous studies of separation measurement in the context of quantum metrology, considering scalar fields, do not apply to fluorescence microscopy. Therefore, the quantum limit of parameter estimation under fluorescence microscopy remained unknown.

In this work, we address this question and determine the quantum limit for measuring the separation of two freely rotating dipoles. We present a method to calculate the QFI involving dipole emitters numerically. We also proposed a near-optimal measurement scheme to estimate the separation of two freely rotating dipoles. In the proposed method, we decouple the radial and azimuthal components of the dipole radiation. The radial component is detected by direct detection for centroid estimation, and the azimuthal component is sent to SLIVER detection for separation estimation. We term this measurement as Polar-SLIVER. A similar configuration was previously proposed for optimal measurement of dipole orientation, but with the radial component entering the SLIVER detection [17].

We numerically investigate the separation estimation of two freely rotating dipoles using different detection methods. We quantify the effects of numerical aperture (NA), detection bandwidth, the number of estimation parameters, background, and misalignment on separation estimation. Our results demonstrate that the proposed detection scheme, Polar-SLIVER, outperforms other methods, especially at high NA. Moreover, we observe that under the ideal condition with no background or imperfections, Polar-SLIVER achieves near-quantum-limited precision on separation estimation.

## 2. Theory

### 2.1. Detection methods

We consider five detection methods based on a combination of direct detection and SLIVER to estimate the separation of two freely rotating dipoles [Fig.1(a)]. The first measurement corresponds to the conventional detection method for most fluorescence microscopes, termed direct detection (DD). The second method uses SLIVER detection, which consists of a Mach-Zehnder interferometer with one arm inverting the electric field in *xy* [Fig.1(e)]. In practice, image inversion can be achieved by placing a dove prism in each arm of the interferometer. One dove prism inverts the input field in *x*, while the other inverts it in *y* [17, 36].

**Fig. 1.**
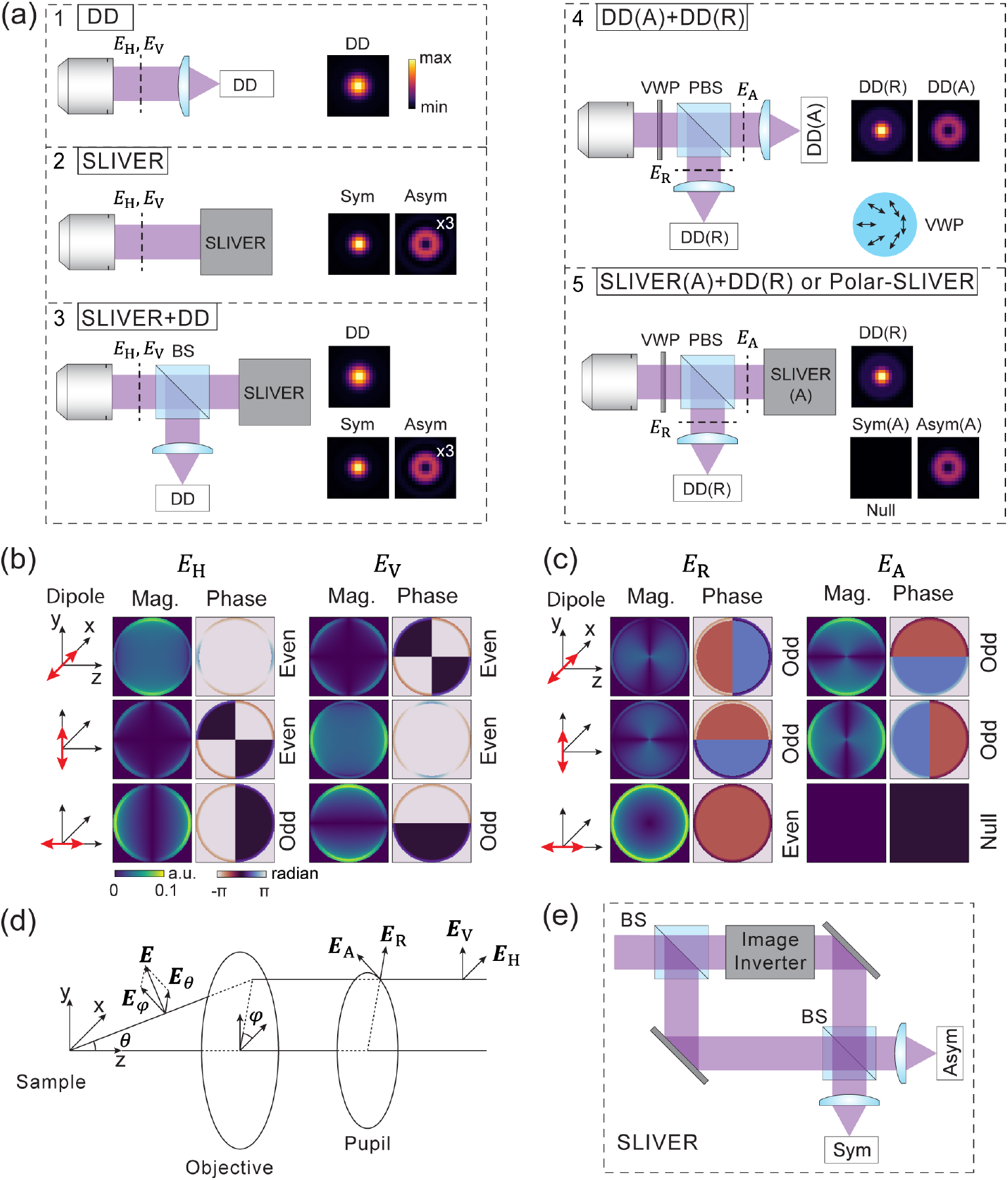
Illustration of five different detection methods for measuring the separation of two freely rotating dipoles. (a) Schematic drawings of the five detection methods considered in this work and their corresponding PSFs. ×*N*: the image contrast is increased by *N* times. DD: direct detection, SLIVER: SLIVER detection, H: horizontal, V: vertical, R: radial, A: azimuthal, Sym: symmetric, Asym: antisymmetric, BS: beam splitter, PBS: polarized beam splitter, Mag: magnitude, VWP: vortex wave plate. A cartoon of VWP illustrates the spatial variant fast axis. The intensity of PSF images is scaled from minimum to maximum values within each detection condition. (b) Pupil magnitudes and phases for horizontal and vertical components of the dipole radiation from dipoles oscillating along *x, y* and *z* axes. a.u.: arbitrary unit. (c) Pupil magnitudes and phases for the radial and azimuthal components of the dipole radiation from a dipole oscillating along *x, y* and *z* axes. (d) Illustration of the coordinate system and definition of the electric fields before and after the objective. See Supplement 1 for the detailed derivation of each electric field component. (e) Schematic drawing of the SLIVER detection.

SLIVER detection decomposes the input field at the pupil plane into its symmetric and antisymmetric components at the output channels. If the input field *h* (*k* _*x*_, *k* _*y*_)is even, such that *h* (*k* _*x*_, *k* _*y*_) = *h* (−*k* _*x*_, −*k* _*y*_), the expected number of photons at the antisymmetric output channel will be zero due to total destructive interference, resulting in a “null output”. Conversely, if the input field is odd, such that *h* (*k* _*x*_, *k* _*y*_) = *h* (−*k* _*x*_, −*k* _*y*_), the symmetric channel results in a null output. Here, we assume *h* (*k* _*x*_, *k* _*y*_) is the input field when a single emitter is positioned at the inversion axis, which coincides with the optical axis of the SLIVER system. If the emitter is offset from the inversion axis to (*x*_*e*_, *y*_*e*_), a tip/tilt phase is introduced to the input field, which can be expressed as *h* (*k* _*x*_, *k* _*y*_)exp (−*ik* _*x*_*x*_*e*_ −*ik* _*y*_ *y*_*e*_). For an even or odd *h* (*k* _*x*_, *k* _*y*_), the tip/tilt phase breaks the field’s symmetry, which results in non-zero photons in the null channel. The high sensitivity of the null channel to a tip/tilt phase allows SLIVER to precisely measure the distance (*d*_*e*_) of a single emitter to the inversion axis, where 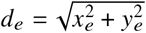. To extend this method for measuring the separation of two emitters, the centroid of the two emitters must be aligned with the SLIVER’s inversion axis. Then the separation measurement is equivalent to measuring 2*d*_*e*_. However, any misalignment of the centroid relative to the inversion axis introduces ambiguity, since both misalignment and separation contribute to photons in the null channel.

As a result, SLIVER by itself cannot accurately measure the centroid position due to the strong correlation between the centroid and separation estimation. To solve this problem, we use a hybrid method similar to the one proposed previously in Ref. [12] and term it SLIVER+DD. In this method, the input field is split equally into two paths. One path is directed to SLIVER, while the other path is directed to a DD measurement. This hybrid measurement method provides three outputs from three channels, one from DD and two from SLIVER. The output from DD is used for estimating the centroid of the two emitters. The fourth method, termed DD(A)+DD(R), decouples the radial and azimuthal components of the input field, which can be realized by adding a vortex wave plate (VWP) after the microscope objective lens. A perfect VWP converts the radially/azimuthally polarized beam (***E***_R_/***E***_A_) into a horizontally/vertically (***E***_H_/***E***_V_) polarized beam with 100% efficiency. With a polarized beam splitter (PBS), the radial and azimuthal components of the dipole radiation can be separated. Each component is collected by a DD measurement. The last method, termed SLIVER(A)+DD(R) (or Polar-SLIVER for short), is our proposed method for measuring the separation of two freely rotating dipoles. In this configuration, the azimuthal component is collected by SLIVER while the radial component is collected by DD. This design is based on the fact that the azimuthal component of dipole radiation is purely odd, independent of the dipole orientation [Fig.1(c)]. Therefore, the symmetric channel of SLIVER ideally will always have total destructive interference at any dipole orientation [Fig.1(a)]. To quantify the contrast of interference, we define the interference visibility for SLIVER as follows,

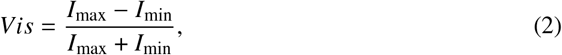

where *I*_max_ and *I*_min_ represent the maximum and minimum output photons from the two channels when imaging a single emitter located at the inversion axis. The visibility achieved by SLIVER(A) is one for detecting a freely rotating dipole, ensuring non-zero FI for estimating the separation of two freely rotating dipoles (Supplement 1).

### 2.2. PSF calculation

The PSFs of the five detection methods shown in Fig. 1 can be calculated as follows,

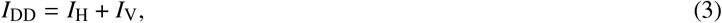

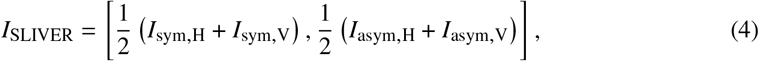

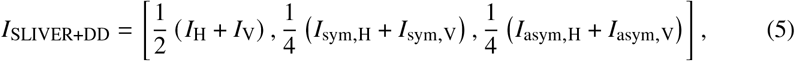

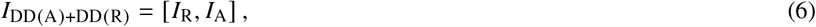

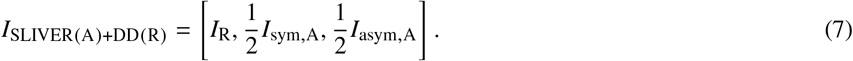

The PSF is defined as the probability distribution of detecting one photon at the image-plane location. The integral of the PSF over the image space is one. For detection methods with multiple channels, the PSF is a vector of the intensity field from all channels. For freely rotating dipoles, the intensity field of the output channels can be calculated from,

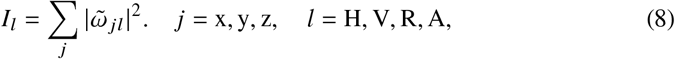

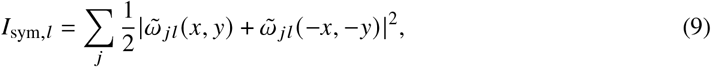

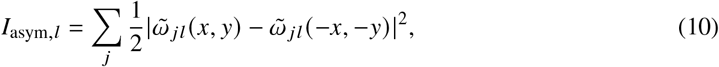

where 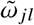 is the amplitude PSF of a dipole oriented in the *j* direction and detected with *l* polarization using DD. The subscript *j* denotes the dipole oriented along *x, y* or *z* direction, while *l* indicates the electric field at the pupil plane in horizontal (H), vertical (V), radial (R), or azimuthal (A) polarization. In this study, we assume that there are no systematic aberrations from the imaging system, such as coma and astigmatism aberrations, induced by optical components. The effect of systematic aberrations on SLIVER detection has been extensively studied in Ref. [37]. Supplement 1 provides detailed derivations of the PSF model for each detection method.

### 2.3. Calculation of the CRB

We model the two incoherent emitters as,

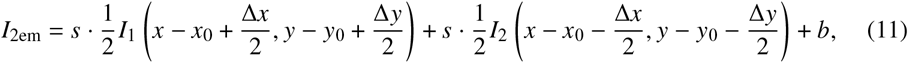

where *I*_1_ and *I*_2_ are PSFs from one of the five detection methods. As two fluorescent molecules of the same type usually have the same photon emission rate under the same excitation power and have the same PSF, we assume that *I*_1_ = *I*_2_, which means that the two emitters are identical in terms of emission rate and shape. The effect of unequal intensities between two emitters has been well studied in previous works [37, 38]. Here, we don’t consider unequal intensities. The displacements Δ*x* = *d* cos *α* and Δ*y* = *d* sin *α* in Eq. (11) represent the separation along the *x* and *y* directions, with *α* being the angle between the *x*-axis and the line defined by two emitters. *s* and *b* denote the total photon and the background photon of the two emitters.

In practice, the intensity field *I*_2em_ is detected by a pixelated camera. Considering the pixelation effect, we define *I*_2em,*q*_ as the integral of the photons from the *q*th pixel of the camera. Assuming that the photon count rate from each pixel follows a Poisson distribution and the pixel values are independent of each other, the CFI matrix is given by [39],

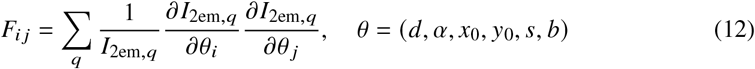

where *θ* denotes the estimation parameters: the centroid position (*x*_0_, *y*_0_), the total photon of two emitters *s*, the separation *d*, the angle *α*, and the background photon per pixel *b*. We then compute the CRB for separation estimation as

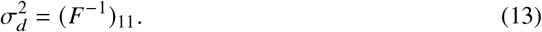

If we consider only the estimation of *d* and assume the other parameters are known, the CRB for estimating *d* is directly given by,

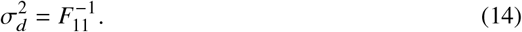

We define

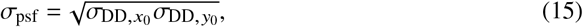

where 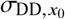 and 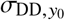 are the precisions for estimating the centroid position (*x*_0_, *y*_0_) of a single freely rotating dipole using DD. As the PSF model in DD is circular symmetric, we have 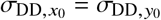.

For DD, when the separation between two emitters approaches infinity, the separation estimation precision *σ*_*d*→∞_ = 2*σ*_psf_ (Supplement 1). In the case of a Gaussian PSF model, 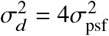 is the QCRB for estimating *d* with an expected photon of one [12].

In the following, unless otherwise specified, we set *s* = 1, *α* = 0, *b* = 0, and *x*_0_ = *y*_0_ = 0. Throughout the text, *σ*_psf_ is always calculated under the condition of *s* = 1 and *b* = 0.

For numerical calculation of the CRB, the two-emitter model *I*_2em_ was generated by simulating a 2D image per channel. The image size was chosen large enough so that the *σ*_psf_ converges. We chose a small pixel size defined by *λ* /8*n*_*a*_, where *λ* is the emission wavelength and *n*_*a*_ the NA of the objective lens. The pixel size is set to a value much smaller than the Nyquist rate to ensure that all features of the PSF patterns are captured. Additionally, even with a small pixel size, the pixel integration needs to be considered. We computed each pixel value as the integral over 2 × 2 up-sampling points within the pixel. Most of the following results were obtained with *λ* = 680 nm and *n*_*a*_ = 1.45, which are typical parameters for oil immersion objectives used in super-resolution imaging of far-red fluorophores. We also incorporated refractive index mismatch (IMM) in the PSF model by setting the refractive indices of the sample medium, coverslip, and immersion oil to 1.33, 1.516, and 1.516, respectively. The corresponding pupil functions are shown in Fig.1(b,c), where the bright rings originate from supercritical angle fluorescence (SAF) caused by IMM [39–42].

### 2.4. Calculation of the QCRB

For the calculation of the QCRB, we note that the electric fields at the pupil plane from dipole radiation are complex vectorial fields. Detailed derivation of the QFI for dipole emitters is shown in Supplement 1; here, we outline the major steps. The QFI for estimating the separation between two freely rotating dipoles is given by

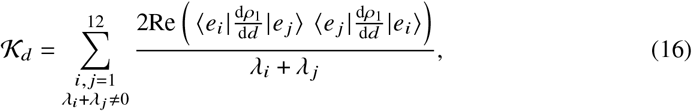

where *ρ*_1_ is the density operator of the one-photon state [17]. For two freely rotating dipoles, the one-photon state can be expressed as,

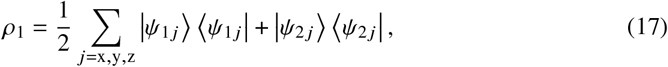

where 1 and 2 denote emitter 1 and emitter 2. Here |*ψ*_1 *j* (2 *j*)_ ⟩ denotes the quantum state of the dipole emitter 1(2) oscillating along the *j* direction, and the vectorial fields are incorporated in the calculation of |*ψ*_1 *j* (2 *j*)_ ⟩. To compute 𝒦_*d*_, *ρ*_1_ is decomposed into its eigenstates |*e*_*i*_⟩ with eigenvalues *λ*_*i*_ as

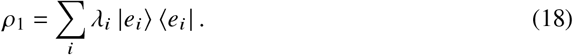

Then, the QCRB for separation estimation is calculated from

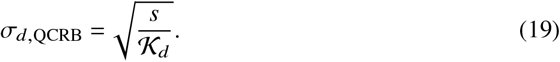

As analytical derivations of the eigenstates are difficult, we compute the QFI numerically (Supplement 1). As *σ*_*d*,QCRB_ is separation (*d*) dependent, we choose its value at infinite separation as a reference to compare the performance of various detection methods,

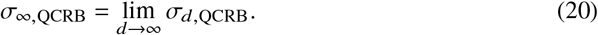

We use this reference as a normalization factor for the following results.

## 3. Results

### 3.1. Precision of separation estimation at different separations

First, we quantified the precision of the separation estimation *σ*_*d*_ for different detection methods as a function of *d* (Fig.2). The two-emitter model *I*_2em_ for each detection method was generated by simulating an image of 150×150 pixels for each channel.

**Fig. 2.**
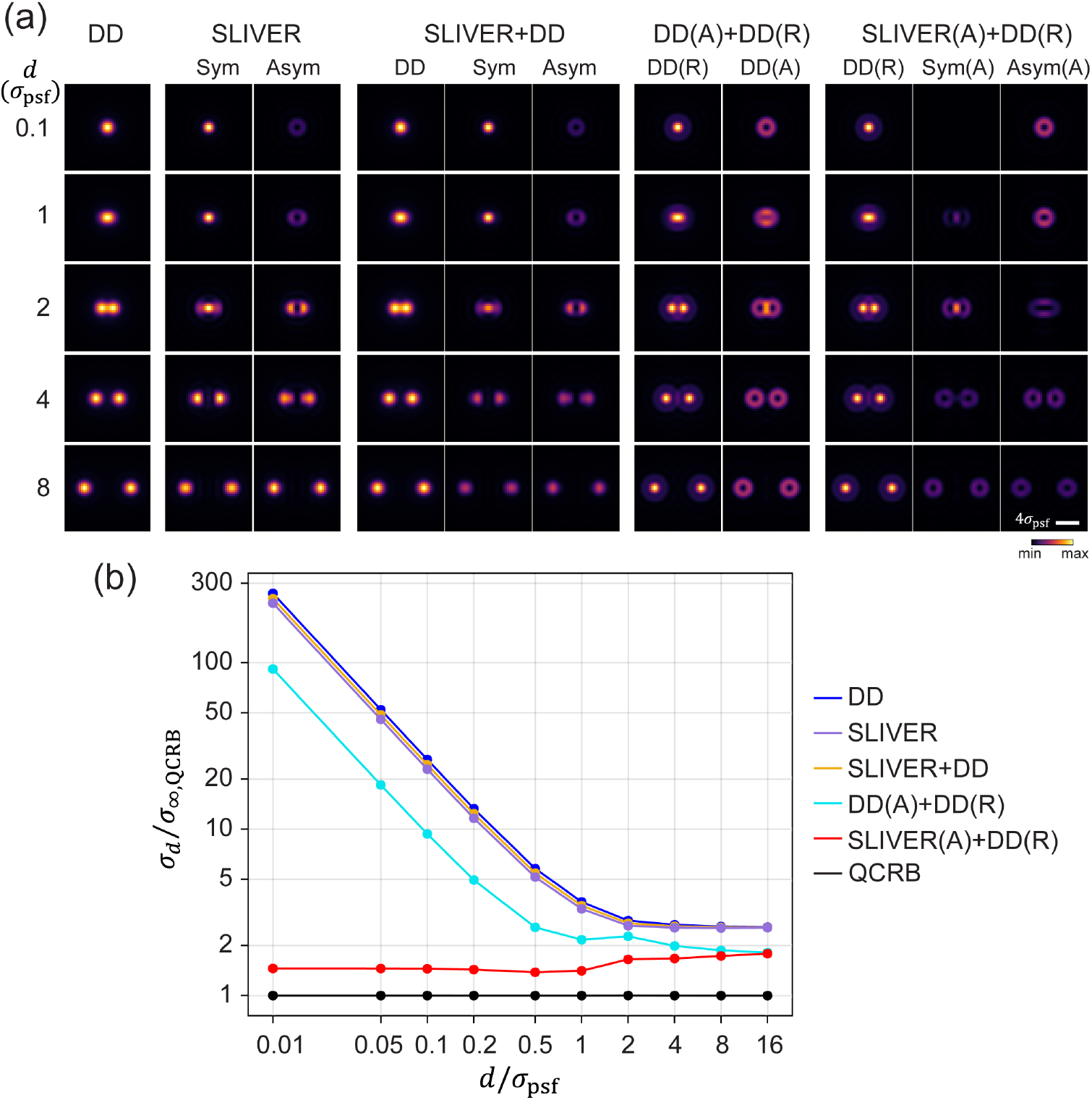
Comparison of separation estimation precision for various detection methods. (a) Two emitter models of different detection methods at various separations. The intensity is scaled from minimum to maximum values within each detection condition. (b) Separation estimation precision *σ*_*d*_ under different detection methods. SLIVER(A)+DD(R) or Polar-SLIVER achieves near-constant precision at all separations, while other methods show large divergence of *σ*_*d*_ as *d* → 0. *σ*_∞,QCRB_ and *σ*_psf_ are defined in the main text.

We found that the precision of separation estimation is comparable across DD, SLIVER, and SLIVER+DD methods at all separations [Fig.2(b)]. SLIVER detection is slightly better than SLIVER+DD, suggesting that SLIVER detection alone is sufficient for estimating separations. However, this method is not practical as it requires prior knowledge of the centroid of two emitters. Furthermore, for measuring the separation of two freely rotating dipoles with a high-NA objective, SLIVER offers little advantage over DD. This finding is in sharp contrast with the previous conclusion drawn from a scalar PSF model, such as a Gaussian PSF model [12].

To understand this difference, we note that the PSF of a dipole emitter is described by vectorial diffraction theory, where its PSF cannot be simply calculated from the Fourier transform of a scalar pupil function, but rather an incoherent sum of the contributions from all polarized fields (Supplement 1). As a result, the radiation from a freely rotating dipole can be decomposed into six polarized fields: three in horizontal polarization (*E*_H_) and three in vertical polarization (*E*_V_), corresponding to dipoles oscillating along *x, y* and *z* directions [Fig.1(b)]. The incoherent sum of those six polarized fields mixes even and odd fields, preventing both the symmetric and antisymmetric channels in SLIVER from achieving total constructive or destructive interference. This results in a significant reduction in interference visibility, which is detrimental to separation estimation, especially at *d* ≪ *σ*_psf_. Even a 0.1% decrease in visibility leads to a rapid divergence in *σ*_*d*_ at small separations, as shown in Fig.3.

**Fig. 3.**
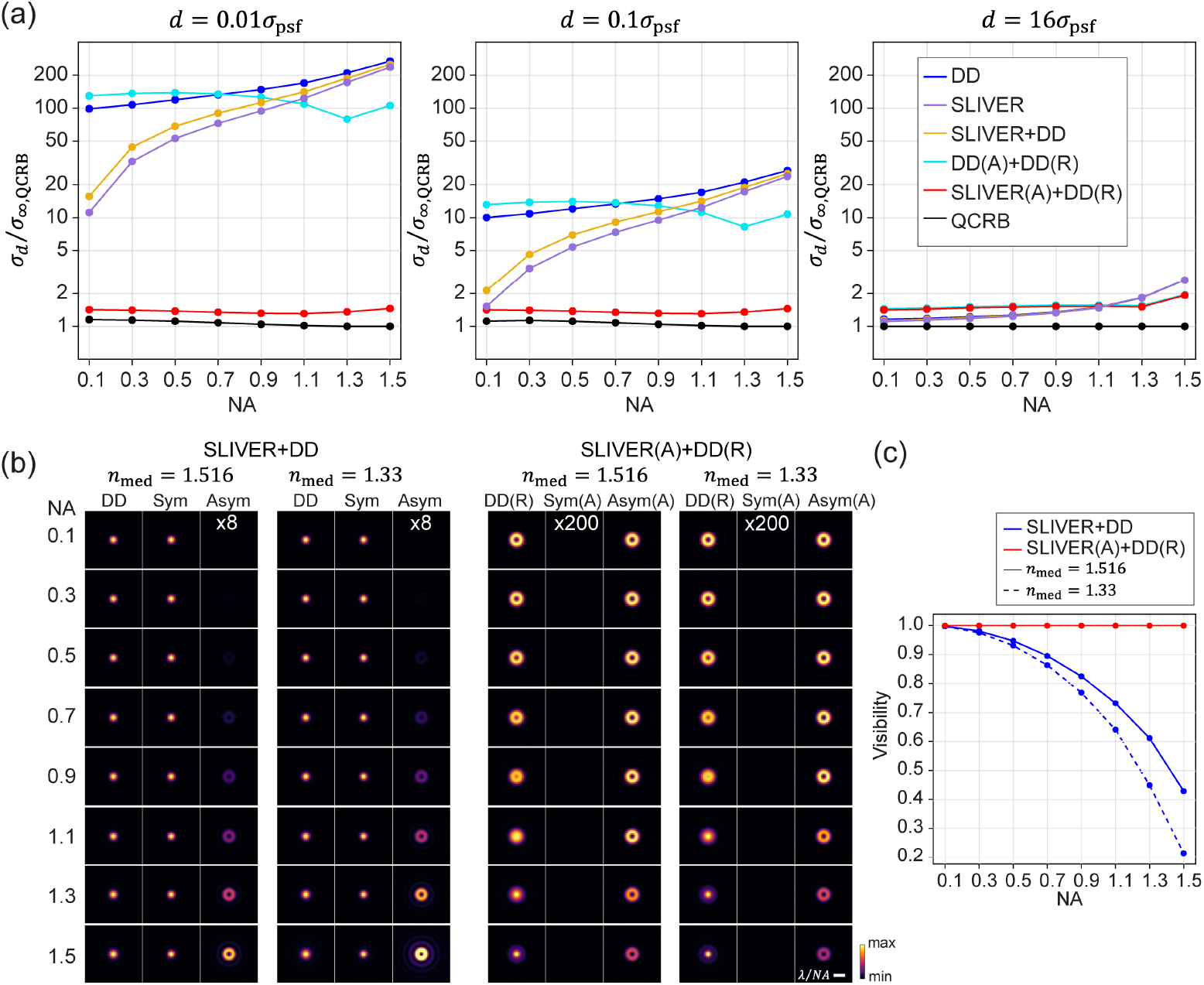
Effect of numerical aperture on separation estimation precision. (a) Comparison of *σ*_*d*_ versus numerical aperture (NA) for different detection methods. Results at separations of 0.01*σ*_psf_, 0.1*σ*_psf_, and 16*σ*_psf_ are shown. The results show that SLIVER(A)+DD(R) or Polar-SLIVER achieves a near-constant value of *σ*_*d*_ for all conditions, while other methods exhibit different levels of divergence in *σ*_*d*_ at small separations. Simulation parameters are: *λ* = 680 nm, refractive index of sample medium *n*_med_ = 1.33, NA at 0.1 to 1.5 with a step size of 0.2, and the rest parameters are the same as those defined in Fig.1. (b) PSFs of different NAs from SLIVER+DD and SLIVER(A)+DD(R). PSFs of index-matched (*n*_med_ = 1.516) and index-mismatched (*n*_med_ = 1.33) conditions are shown. For SLIVER+DD, the signal from the anti-symmetric channel increases with NA, while for SLIVER(A)+DD(R), the symmetric channel remains dark at all NAs. (c) Comparison of visibilities from SLIVER+DD and SLIVER(A)+DD(R). Visibility of SLIVER(A)+DD(R) remains one at all NAs. Visibility of SLIVER+DD decreases with NA, and the decrease is faster in the case of index mismatching.

To recover the estimation precision for freely rotating dipole emitters, we propose to decouple the radially (*E*_R_) and azimuthally (*E*_A_) polarized fields of the dipole radiation. To this end, we introduce a VWP after the objective and replace the beam splitter (BS) with a PBS. Note that each polarization (R and A) has three field components from three orthogonally orientated dipoles [Fig.1(c)]. We identified that azimuthally polarized fields exhibit purely odd symmetry, which is optimal for SLIVER detection. This configuration, Polar-SLIVER, allows for total constructive interference of the field in the A component, thereby achieving a visibility of one. Fig.2(b) shows that Polar-SLIVER recovers the precision of separation estimation and achieves *σ*_*d*_ near the QCRB for any separation. Furthermore, we note that *σ*_*d*_ as *d* → 0 is even smaller than the values as *d* → ∞ (Fig. S1). We show that this is caused by dipole radiation, where the PSF is the incoherent sum of fields from multiple dipole orientations. This summation leads to different values at the limits of *d* →0 and *d* → ∞ (Supplement 1). In terms of FI, Polar-SLIVER reaches around 50% of the QFI at small separations, while the FI of other methods is up to 15-25% of the QFI and drops to zero as *d* → 0 (Fig.S1 and Fig.S2). The missing 50% of the FI in Polar-SLIVER is due to contributions of *z*-oriented dipole (Supplement 1).

To investigate the effect by simply decoupling *E*_R_ and *E*_A_, we calculated *σ*_*d*_ under direct detection of *E*_R_ and *E*_A_, DD(A)+DD(R). We found that although *σ*_*d*_ of DD(A)+DD(R) still diverges as *d* → 0, the divergence is about two times smaller than that of DD. This is because the sharper features of the PSFs produced by radially and azimuthally polarized fields [Fig.2(a)] contains more information about the separation, which improves its estimation precision. Furthermore, as *d* → ∞, *σ*_*d*_ of DD(A)+DD(R) and Polar-SLIVER converge to the same value. This is because their corresponding FI is equal at *d* → ∞(Supplement 1). We observed a subtle feature in the variation of *σ*_*d*_ versus *d* for both Polar-SLIVER and DD(A)+DD(R) within a separation of 0.5*σ*_psf_ to 2*σ*_psf_. This behavior can be attributed to changes in the FI within each channel of the respective detection methods (Fig. S2). For Polar-SLIVER, the FI of SLIVER(A) is maximum at *d* = 0; however, the inclusion of the DD(R) channel shifts the maximum of FI to *d* ≈ 0.7*σ*_psf_. For DD(A)+DD(R), the FI of both channels exhibits a local peak at *d* ≈ 0.9*σ*_psf_, leading to a local maximum at the same location in the total FI. In contrast, detection methods that do not employ polarization sorting in A and R by the VWP, such as SLIVER+DD, show a smoother change in FI with respect to *d* for both dipole and Gaussian PSFs. This indicated that PSFs from radial and azimuthal polarizations exhibit sharper features.

### 3.2. Precision of separation estimation at different numerical apertures

We next investigate the effect of NA on separation estimation [Fig.3(a)]. We found that *σ*_*d*_ from the three methods without polarization sorting (no VWP) increase with NA. At low NA (NA<1.1), SLIVER and SLIVER+DD outperform both DD and DD(A)+DD(R). This is because, as NA decreases, the contribution from the *z*-oriented dipoles diminishes when most of its radiation is lost at small NAs. Therefore, the radiation collected from a freely rotating dipole predominantly generates even fields, which enhances the interference visibility [Fig. 3(c)] and improves the precision of the estimator. However, even when NA is reduced to 0.1 and the visibility increases to 99.8% [Fig.3(c)], *σ*_*d*_ from SLIVER and SLIVER+DD still diverges as *d* → 0 [Fig.3(a)], increasing by 10 -fold from *d* = 16*σ*_psf_ to *d* = 0.01*σ*_psf_. At high NA (NA>1.1), both SLIVER and SLIVER+DD perform worse than DD(A)+DD(R) and approach the performance of DD. We also found that IMM (*n*_med_ = 1.33) will further reduce the visibility of SLIVER and SLIVER+DD as NA increases [Fig.3(b,c)]. However, this reduction in visibility has a small influence on the separation estimation [Fig.3(a) and Fig. S3].

We show that polarization sorting with a VWP improves separation estimation precision at high NA for DD(A)+DD(R). For Polar-SLIVER, as the visibility remains constant and equal to one at all NAs [Fig.3(c)], *σ*_*d*_ does not diverge as *d* → 0 [Fig.3(a)]. In the case of IMM, *σ*_*d*_ for DD(A)+DD(R) is further reduced at high NA. However, when NA increases to ∼1.3, the FI for estimating *d* starts to decrease for DD(A)+DD(R) at all separations and for Polar-SLIVER at *d* → ∞[Fig. S1(b)]. This decrease in FI results in a corresponding increase in *σ*_*d*_ [Fig. 3(a) and Fig. S1(a)]. We also found that as *d* → ∞, the detection methods with and without VWP converge to different points [Fig.3(a)]. This difference is because the PSFs without VWP result from horizontally and vertically polarized fields [Fig.1(b)], while the PSFs with VWP result from azimuthally and radially polarized fields [Fig. 1(c)]. This leads to different FI for estimating *d* as*d* → ∞ (Supplement 1).

### 3.3. Practical considerations

#### 3.3.1. Effect of detection bandwidth

The above calculations consider an ideal VWP, which can convert a radially/azimuthally polarized field to a horizontally/vertically polarized field at 100% efficiency. However, fluorescence emission often has a broad spectrum. To achieve 100% conversion efficiency, a single VWP has to be optimized for a broad spectral range. To our knowledge, broadband VWP is more complicated in fabrication and has lower transmission efficiency [43]. Here we consider a commonly used commercial VWP (e.g. WPV10L-633, Thorlabs) that is only optimized for a single wavelength. This VWP is fabricated from nematic liquid crystal (LC), with LC molecules aligned azimuthally to create a half-wave plate with spatially varied fast axis [44]. We simulate the VWP based on the dispersion of nematic LC, where the phase delay between the extraordinary and ordinary polarizations is given by *δ* = 2*π* (*n*_e_ − *n*_o_) *t* /*λ*, with *t* being the thickness of the VWP and *n*_*e*_ and *n*_*o*_ being the refractive indices of the corresponding polarizations. This VWP can only be optimized for a single wavelength that satisfies the half-wave delay condition, *δ* = *π*, ensuring 100% conversion efficiency. However, the conversion efficiency of the VWP decreases as the detection bandwidth (Δ*λ*), defined by the emission filter, increases. With partial conversion, the fields entering the SLIVER detection arm will not be purely azimuthally polarized, preventing total destructive interference in the symmetric channel [Fig.4(a)]. Consequently, the visibility reduces with increasing Δ*λ*, leading to a divergence of *σ*_*d*_ as *d* → 0 [Fig.4(b,c)]. Due to the larger dispersion of nematic LCs at shorter wavelengths, the visibility decreases faster as Δ*λ* increases, resulting in a larger divergence of *σ*_*d*_ at shorter wavelengths (Fig. S4). For super-resolution imaging, the emission filter typically has a bandwidth of 30-50 nm to achieve a high signal-to-noise ratio. Within this bandwidth range, the divergence is relatively small compared to the other four methods [Fig.2(b)].

**Fig. 4.**
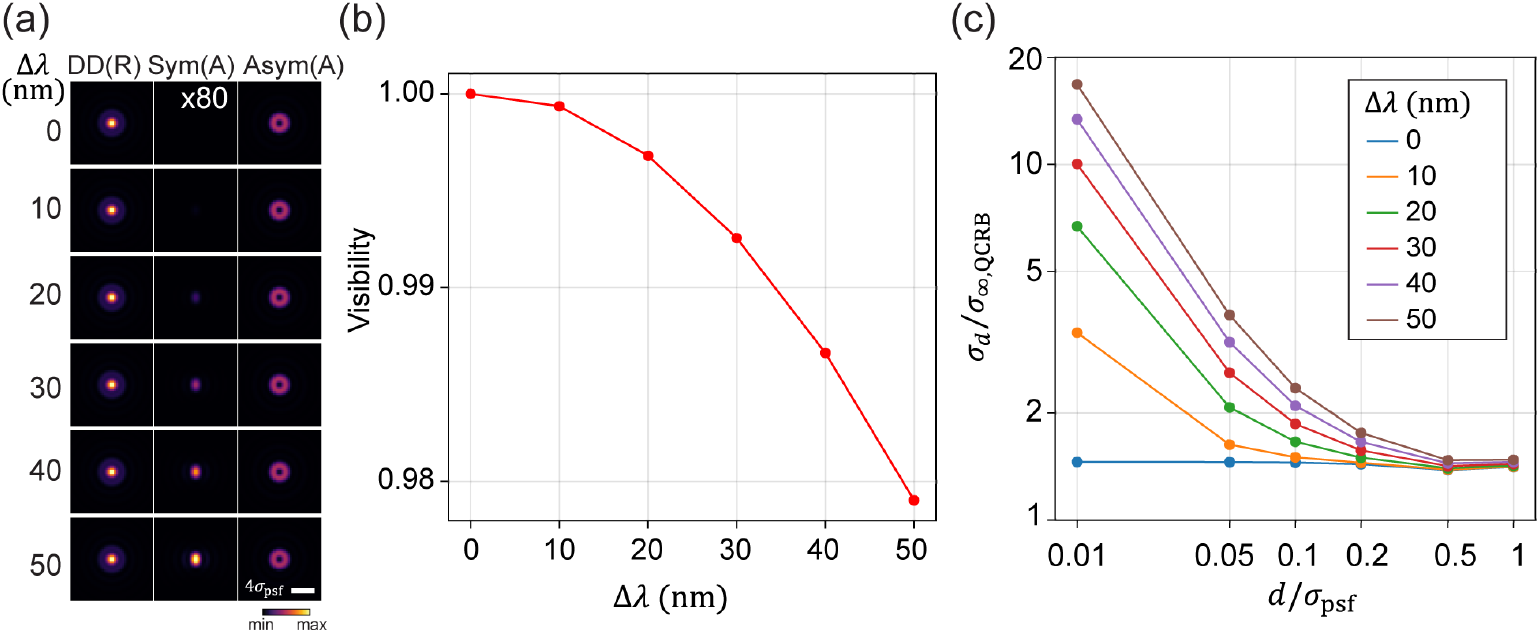
Effect of detection bandwidth on separation estimation for Polar-SLIVER. (a) PSFs of Polar-SLIVER at various detection bandwidths, Δ*λ*. The signal from the symmetric channel increases with detection bandwidth, leading to a reduction of the visibility. (b) Visibility versus Δ*λ*. (c) *σ*_*d*_ versus separation at various Δ*λ. σ*_*d*_ starts to diverge as *d* → 0 with a non-zero detection bandwidth. Simulation parameters are: *n*_*a*_ = 1.45, *λ* = 680 nm, *n*_med_ = 1.33, the VWP is optimized at 680 nm.

#### 3.3.2. Effect of the number of estimation parameters

The above results were obtained by considering the separation *d* as the only estimation parameter, assuming other parameters are known. We next examine the estimation precision of *d* when estimating all parameters together (full-parameter estimation). The parameters were defined in Eq. (11). Here, we focus on the Polar-SLIVER and DD methods. We found that *σ*_*d*_ has negligible variations between full-parameter (six parameters) and one-parameter estimations for both detection methods under the ideal condition [Fig. 5(a)]. We next investigate the impact of the number of estimation parameters when introducing a non-zero background photon per pixel or a slight misalignment of the centroid positions of two emitters (refer to sections 3.3.3 and 3.3.4 for further discussion on background and misalignment). We found that under the condition of a non-zero background, the number of estimation parameters also has little effect on *σ*_*d*_ for both Polar-SLIVER and DD [Fig. 5(b)]. However, for Polar-SLIVER, when the centroid of two emitters is misaligned with the inversion axis of SLIVER(A), *σ*_*d*_ is smaller when estimating *d* only [Fig. 5(c)]. For DD, since the separation estimation is independent of the centroid position, an offset of the centroid position does not affect the estimation of *d* in both one-parameter and full-parameter estimations. Furthermore, as the PSF in Polar-SLIVER is circular symmetric, the angle *α* has little effect on the separation estimation in full-parameter estimation [Fig. S5(b,c)], and *σ*_*d*_ is the same as the one from one-parameter estimation [Fig. 2(b)]. We noticed that the angle estimation precision *σ*_*α*_ diverges quickly as *d* → 0, which implies that Polar-SLIVER is not optimal for angle estimation [Fig. S5(d,e)]. However, compared to DD, Polar-SLIVER still performs better in angle estimation [Fig. S6(b) and Fig. S7(b)].

**Fig. 5.**
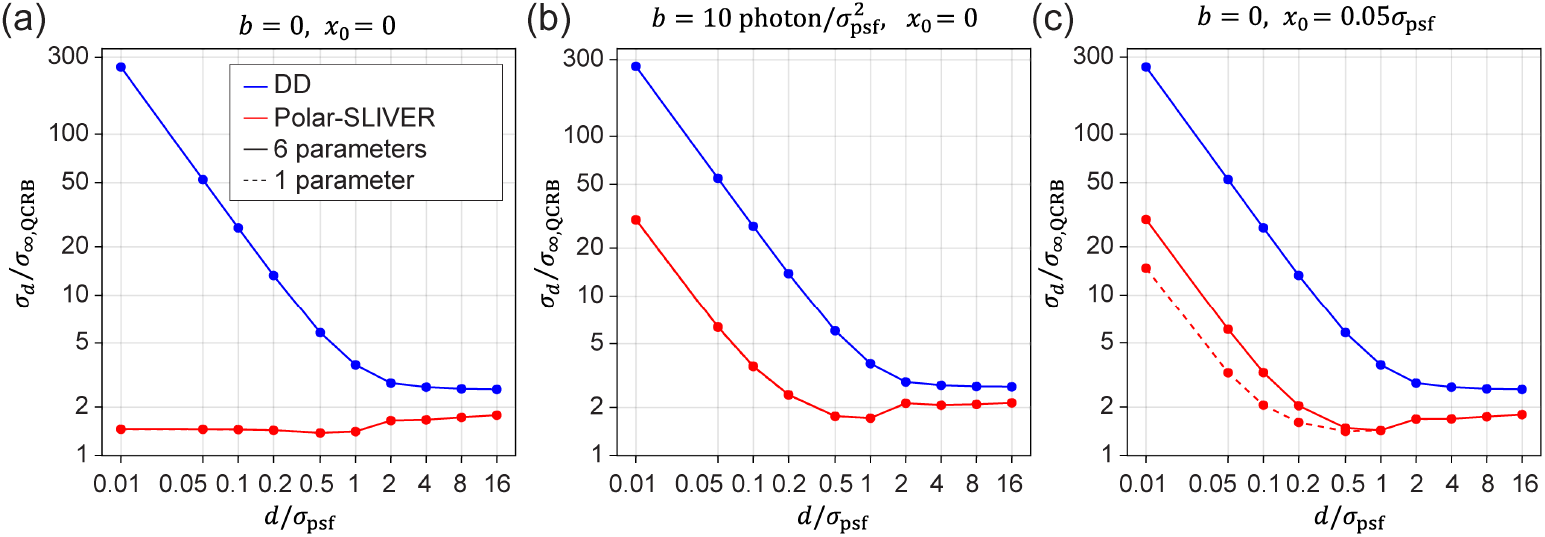
Effect of the number of estimation parameters on separation estimation. Precisions of separation estimation when estimating *d* only (solid lines) and when estimating all six parameters together (dash lines), using DD or Polar-SLIVER, are shown. (a) Results under the ideal condition. There is no obvious difference between one and full parameter estimation. (b) Results under the condition of non-zero background: 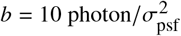, total photon *s* = 10000. *σ*_*d*_ of Polar-SLIVER starts to diverge as *d* → 0. There is also no obvious difference between one and full parameter estimation. (c) Results under the condition of misalignment: *x*_0_ = 0.05*σ*_psf_. *σ*_*d*_ of Polar-SLIVER starts to diverge as *d* →0. One parameter estimation is slightly better for Polar-SLIVER, indicating a strong correlation between centroid and separation estimation. Simulation parameters are: *n*_*a*_ = 1.45, *λ* = 680 nm, *n*_med_ = 1.33, total photon *s* = 1 for (a) and (c), VWP is ideal.

### 3.3.3. Effect of background on separation estimation

In fluorescence imaging, background is often inevitable. Background mainly comes from out-of-focus fluorophores, sample autofluorescence, and impurities of the coverslip surface. Here, we investigate the impact of a non-zero background on separation estimation. We simulated data with 10,000 total photons per measurement and added a constant background photon per pixel. We define a background unit as photons per 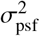, which can be interpreted as photons per pixel when the pixel size is *σ*_psf_. We found that for Polar-SLIVER, even with a background level of 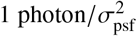, the precision of separation estimation starts to diverge [Fig. 6(a)] as *d* → 0. This divergence continuously increases with increasing background. However, within a common background range 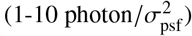 at low labeling density, Polar-SLIVER still outperforms DD by a factor of 9 as *d* → 0. We noticed that the background has minimal impact on separation estimation using DD. This is also true for angle and centroid estimation, where background has less effect on DD [Fig. S6(b-d)]. For estimations of total photon and background photon, the background has a similar effect on both DD and Polar-SLIVER [Fig. S6(e,f)]. Additionally, adding a detection bandwidth of 30 nm further increases *σ*_*d*_, however, the increase is less at higher background levels [Fig. 6(b)].

**Fig. 6.**
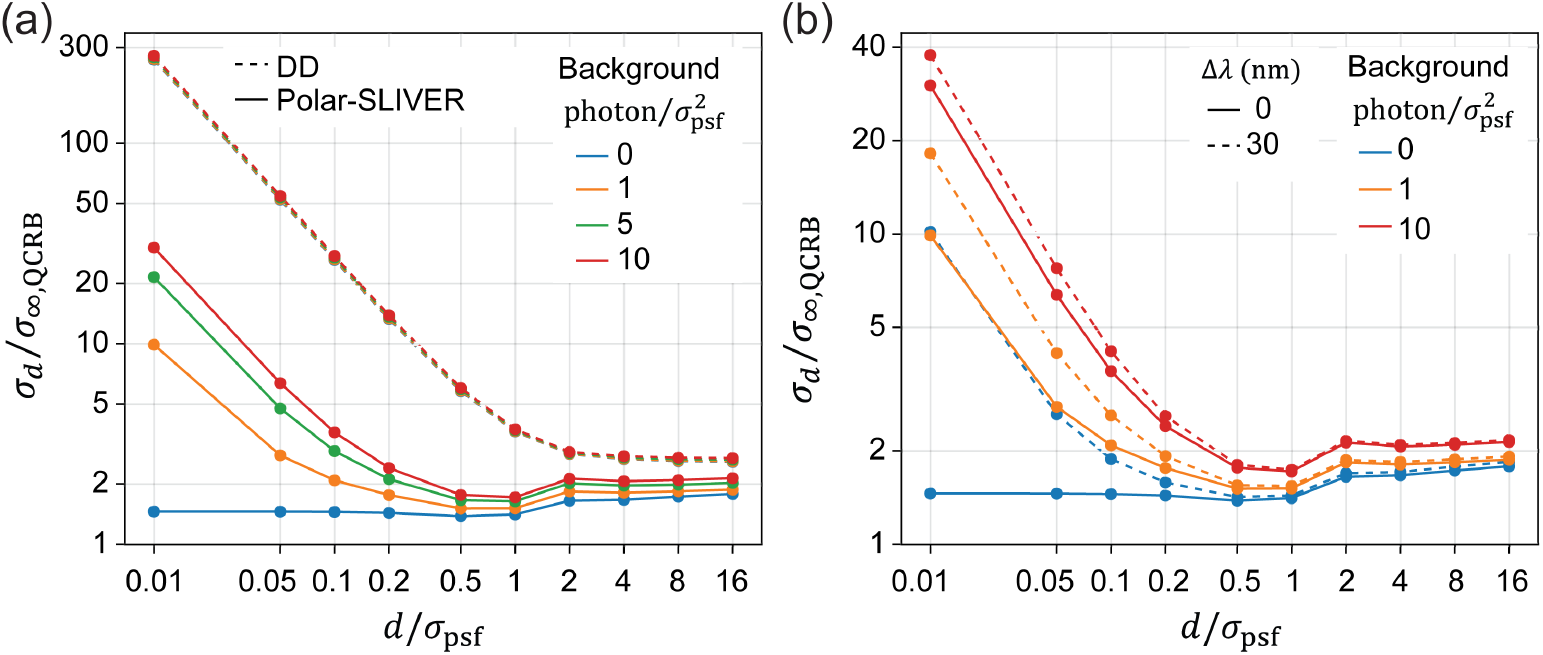
Effect of background on separation estimation. (a) Comparison of separation estimation of DD and Polar-SLIVER at different background levels. *σ*_*d*_ of Polar-SLIVER starts to diverge as *d* → 0 with non-zero background. Simulation parameters are: *n*_*a*_ = 1.45, *λ* = 680 nm, *n*_med_ = 1.33, total photon *s* = 10000, VWP is ideal. (b) Separation estimation precision of Polar-SLIVER at different background levels and detection bandwidths. Both non-zero background and detection bandwidth will induce divergence of *σ*_*d*_ as *d* →0. However, at a higher background level, the effect of detection bandwidth becomes smaller. Simulation parameters are: *n*_*a*_ = 1.45, *λ* = 680 nm, *n*_med_ = 1.33, total photon *s* = 10000, VWP is optimized at 680 nm. Precisions were calculated under full parameter estimation.

We also examined the performance of Polar-SLIVER at different levels of signal-to-background ratio (SBR) (Fig. S8). We define SBR as

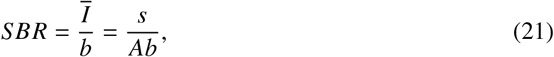

where *Ī* represents the average intensity, calculated from the total photon *s* divided by an area *A*. Here *A* is set to 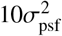, encompassing the majority of the PSF. Our results showed that with an ideal VWP and no misalignment, Polar-SLIVER is always >2 times better than DD in separation estimation. When accounting for an imperfect VWP and a detection band width of 30 nm, Polar-SLIVER maintained its performance at low SBR, demonstrating the dominant influence of SBR in such conditions. However, at high SBR levels, the impact of the imperfect VWP became more pronounced. Furthermore, misalignment of the centroid position has a larger effect on Polar-SLIVER (refer to section 3.3.4 for further discussion on misalignment). At a misalignment of 0.2*σ*_psf_, the performance of Polar-SLIVER reduces to that of DD(A)+DD(R), while at a misalignment of 0.5*σ*_psf_, it reduces to that of DD. Assuming that misalignment is primarily determined by the estimation precision of the centroid position, 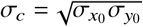, Polar-SLIVER continues to outperform other methods when SBR>1.

### 3.3.4. Effect of misalignment on separation estimation

Lastly, we examine the effect of misalignment between the centroid of two emitters and the inversion axis of SLIVER(A). For Polar-SLIVER, the FI for separation estimation drops to zero as *d* →0 with even a small misalignment of the centroid position. We introduce a misalignment by adding an offset to the centroid position along the x-axis (*x*_0_). Figure 7(a) shows that at a misalignment of 0.01 to 0.1*σ*_psf_, *σ*_*d*_ increases by 3.5 to 35 times at *d* = 0.01*σ*_psf_. However, for DD, as there is no image inversion, the centroid position has no impact on the separation estimation [Fig. 7(a)]. This is also true for estimating other parameters using DD [Fig. S7(b-f)]. For Polar-SLIVER, misalignment will also reduce the precision of angle estimation, but slightly improve the precision of centroid estimation [Fig. S7(b-d)]. Next, we study the condition when combining the effects of detection bandwidth, background, and misalignment. We find that with a detection bandwidth of 30 nm, the addition of a misalignment of 0.05*σ*_psf_ further increases *σ*_*d*_ by 2-4 fold under different background levels. This indicates that misalignment has a more dominant effect on separation estimation compared to detection bandwidth and background. In practice, the precision of centroid estimation is around 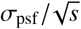, where *s* is the total photon count of two emitters. As discussed above, if the misalignment is mainly due to the centroid estimation precision, to maintain the misalignment within 0.05*σ*_psf_, *s >* 1000 is required (Fig. S6-S7).

**Fig. 7.**
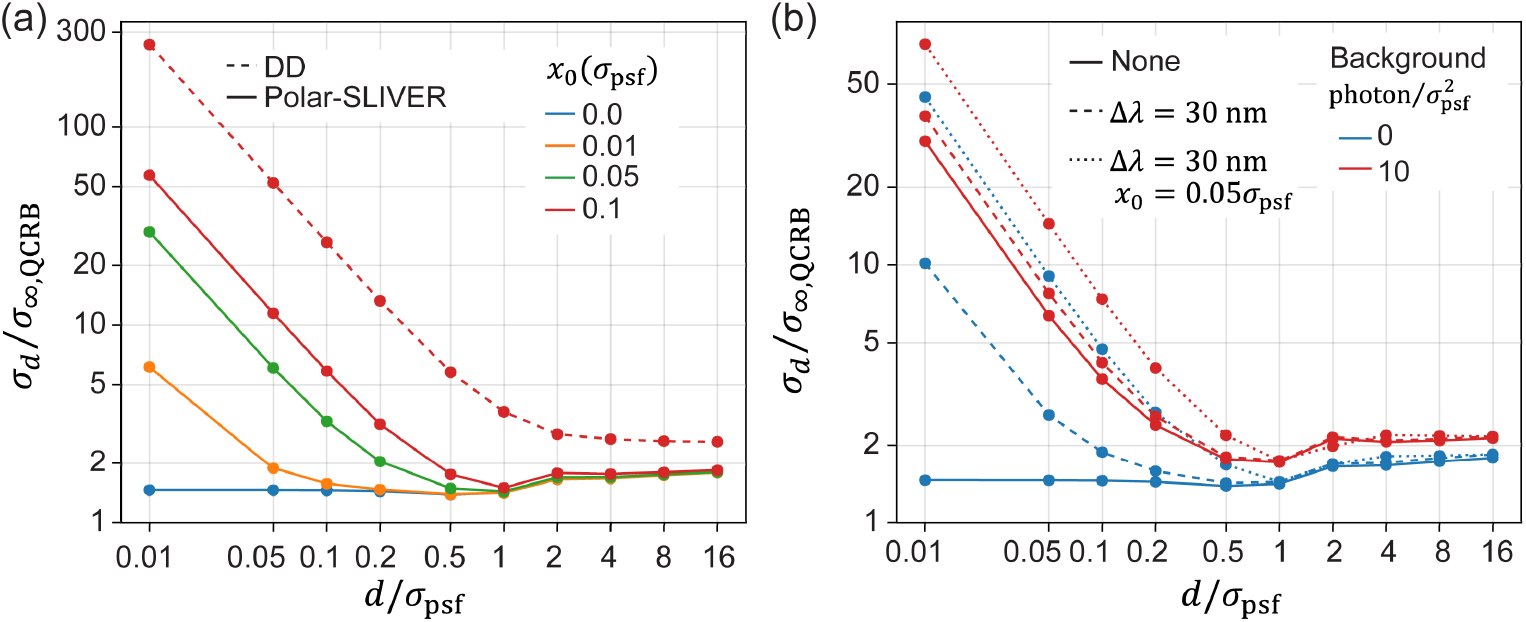
Effect of misalignment on separation estimation. (a) Comparison of separation estimation of DD and Polar-SLIVER at different amounts of misalignment. Misalignment is introduced by adding an offset to the centroid position along the x-axis (*x*_0_). *σ*_*d*_ of Polar-SLIVER starts to diverge as *d* →0 with non-zero misalignment. Simulation parameters are: *n*_*a*_ = 1.45, *λ* = 680 nm, *n*_med_ = 1.33, total photon *s* = 1, background *b* = 0, VWP is ideal. (b) Separation estimation precision of Polar-SLIVER at different background levels, under the conditions of zero detection bandwidth and zero misalignment (solid line), a detection bandwidth of 30 nm and zero misalignment (dash line), and a detection bandwidth of 30 nm plus a misalignment of 0.05*σ*_psf_ (dotted line). Non-zero background, detection bandwidth, and misalignment will all induce divergence of *σ*_*d*_ as *d* → 0. However, at a higher background level (red lines), the effect of misalignment is greater than that of detection bandwidth. Simulation parameters are: *n*_*a*_ = 1.45, *n*_med_ = 1.33, total photon *s* = 10000, VWP is optimized at 680 nm. Precisions were calculated under full parameter estimation.

## 4. Discussion and Conclusion

In this work, we present a numerical method for calculating the QFI of estimating the separation between two dipole emitters and introduce a novel detection method, Polar-SLIVER, that approaches quantum-limited precision. We evaluated separation estimation using five different detection methods and demonstrated that Polar-SLIVER outperforms the others. Under the ideal condition, Polar-SLIVER achieves near quantum-limited precision in separation estimation. We also examined the impact of numerical aperture, detection bandwidth, the number of estimation parameters, background, and misalignment on separation estimation. Our findings indicate that, even with these practical considerations, Polar-SLIVER continues to outperform the other methods.

To experimentally realize Polar-SLIVER, several aspects should be considered. First, Polar-SLIVER relies on a device that can convert radial/azimuthal polarization to horizontal/vertical polarization. This conversion falls into the category of spin-angular-momentum (SAM) to orbital-angular-momentum (OAM) conversion of light [45, 46]. Many techniques have been developed to achieve SAM-OAM conversion. Future development will include exploring the optimal SAM-OAM conversion method for the experimental implementation of Polar-SLIVER. Furthermore, SLIVER(A) requires the centroid of two emitters to be aligned with the inversion axis, which limits the measurement to one dimer molecule at a time. To achieve high-throughput imaging, Polar-SLIVER can be combined with high-speed single particle tracking for fast repositioning of the centroid. This combination will also allow Polar-SLIVER to track moving molecules: the estimated centroid position is sent to a feedback system, which re-centers the centroid position with the inversion axis of SLIVER(A).

Additionally, the interferometer in SLIVER(A) should be stabilized at zero phase delay. Existing phase stabilization methods achieve a precision of 1 degree [47, 48], which is sufficient for the proposed method [Fig. S9(a)]. Intensity split ratio between the two interference arms is another consideration factor, but its effect is relatively small [Fig. S9(b)]. A split ratio of 1:0.95 is already adequate for the proposed method.

Furthermore, although we used a small pixel size in most of our simulations, we found that pixel size has minimal impact on separation estimation using Polar-SLIVER (Fig. S10). At varying SBR levels, using a pixel size larger than the Nyquist rate results in a 15-30% decrease in separation estimation precision. Therefore, we recommend choosing a pixel size close to the Nyquist rate. To exploit the full potential of Polar-SLIVER, techniques that minimize background fluorescence are also critical for actual implementation.

Aberrations in the imaging system also impact the precision of separation estimation. Previous work by Schodt et al. examined the effect of various aberrations on a SLIVER system using a scalar PSF model [37]. Their findings are equally applicable to Polar-SLIVER systems employing a vectorial PSF model. Here, we give a brief summary of the effect from aberrations. Aberrations with even symmetry in their PSF patterns, such as spherical and astigmatism aberrations, have little effect on Polar-SLIVER. However, asymmetric aberrations, such as coma and trefoil, reduce the visibility of SLIVER(A), thereby affecting its performance. Furthermore, independent aberrations between the two arms of the interferometer can degrade the performance of Polar-SLIVER, and this issue can arise from any type of aberration. In practice, systematic aberrations can be minimized by using a wavefront control device (e.g., deformable mirror) in each arm of the interferometer [35].

Our proposed method of decoupling the radial and azimuthal components of dipole radiation can also benefit other modal imaging techniques, such as SPADE. To achieve high estimation precision, SPADE requires near-zero crosstalk between orthogonal optical modes. However, without VWP, the input fields from a freely rotating dipole can be approximated as a mixture of the first three Hermite-Gaussian (HG) modes, substantially reducing SPADE’s sensitivity in separation estimation. In contrast, input fields from azimuthal polarization contribute minimally to HG_00_, making the HG_00_ channel highly sensitive to emitters’ separations. It has been shown that SPLICE does not achieve QCRB, even for a Gaussian PSF [24]. Furthermore, SPLICE is not applicable to dipole emitters, because no matter which polarization component is used, the output counts from SPLICE will not vanish at zero separation. Similarly, a SLIVER system with a one-dimensional image inverter is also not applicable to dipole emitters.

In this work, we focus on the separation estimation of two emitters. In the case of more than two emitters, the estimation parameters include a set of pairwise distances. It has been shown that for a one-dimensional array of emitters, no more than two distances can be measured with non-zero QFI as distances approach zero [49]. This implies that no more than two pairwise distances can be unknown parameters, limiting the number of emitters to two. Alternatively, image moments can be used to fully describe the emitter distribution instead of pairwise distances [50]. Zhou et al. demonstrated that up to the second-order moment can be measured with non-zero QFI as emitters’ distances approach zero [51]. Tsang et al. derived the QCRB for estimating image moments [52] and demonstrated that SPADE can approach the QCRB for image moment estimation [20, 53]. Although these studies are based on Gaussian PSFs, their conclusions can be extended to dipole emitters. We anticipate that a modified SPADE system with a VWP can be used to measure up to the second-order moment of multiple dipole emitters at arbitrary separations.

In summary, we anticipate that Polar-SLIVER will enable distance measurement of dipole emitters to approach the quantum-limited precision and will find broad applications in fluorescence microscopy, where many fluorescence molecules are modeled as dipole emitters. Furthermore, our numerical method of QFI calculation for vector fields can be broadly applied to other estimation problems, such as position estimation, and inspire future exploration of optimal measurement strategies for dipole emitters.

## Data availability

No data were generated or analyzed in the presented research.

## Supplemental document

See Supplement 1 for supporting content.

