## Supplementary File for "Separation estimation of two freely rotating dipole emitters near the quantum limit"

### Supplementary Figures

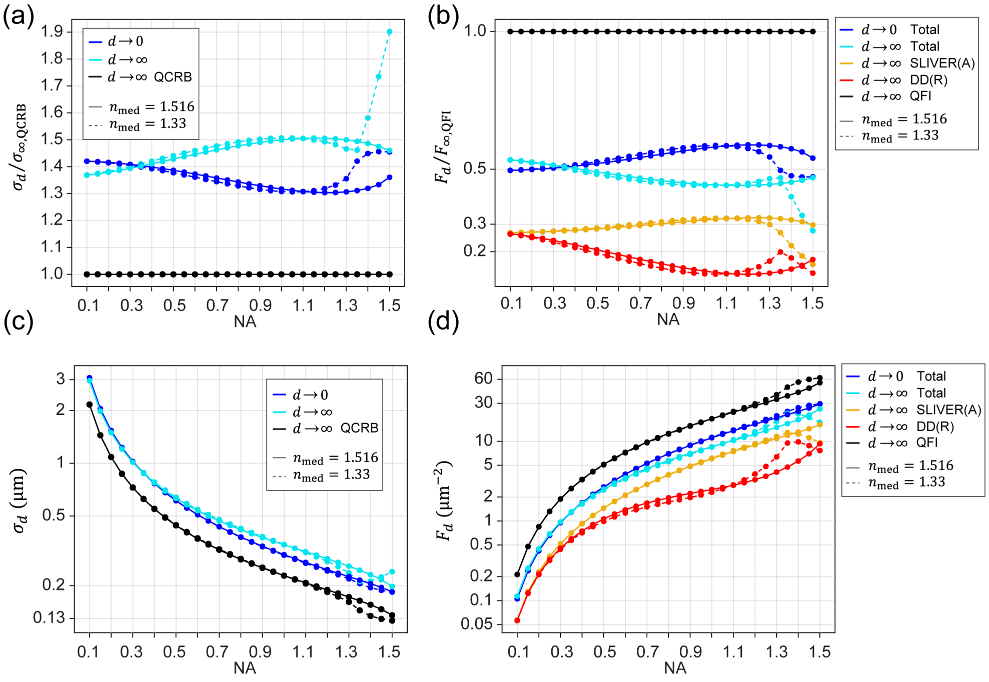


**Fig. S1.** Theoretical precision and Fisher information for separation estimation using SLIVER(A)+DD(R) or Polar-SLIVER. (a) Precision of separation estimation at $d\to0$ and $d\to\infty$. Results under refractive index-matched ($n_{\mathrm{med}}=1.516$) and mismatched ($n_{\mathrm{med}}=1.33$) conditions are shown. (b) Fisher information (FI) for separation estimation $F_d$ at $d\to0$ and $d\to\infty$. Total FI of $d\to0$ is from the symmetric channel of SLIVER(A), while the total FI of $d\to\infty$ is from both SLIVER(A) and DD(R) channels. Results under refractive index-matched and mismatched conditions are shown. $F_{\infty,\mathrm{QFI}}$ is the quantum FI (QFI) for estimating the separation of two freely rotating dipoles as $d\to\infty$. (c) Same as (a), but with a physical unit. $\sigma_d$ decreases with NA. However, index mismatching causes a turning point in $\sigma_d$ of $d\to\infty$ at NA around 1.35. (d) Same as (b), but with a physical unit. We again observe a turning point in $F_d$ of $d\to\infty$ at NA$\sim$1.35 when the index is mismatched. This is because supercritical angle fluorescence (SAF) enhances side lobes of the PSF patterns, reducing the information under the same number of photons. Simulation parameters are: $\lambda=680$ nm, VWP is ideal.

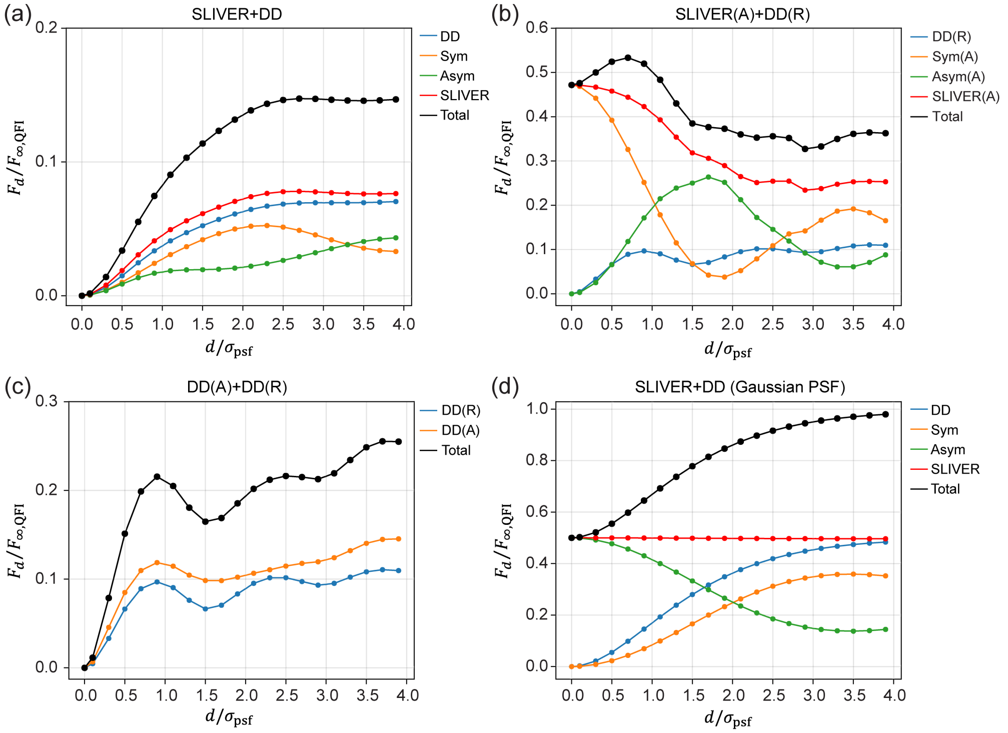


**Fig. S2.** Fisher information for separation estimation versus separation for different detection methods. (a) Fisher information (FI) of SLIVER+DD. SLIVER denotes the sum of FIs from symmetric and antisymmetric channels. Total indicates the FI of all channels. $F_{\infty,\mathrm{QFI}}$ is the quantum FI (QFI) for estimating the separation of two freely rotating dipoles as $d \to \infty$. We observe that the total FI reduces to zero as $d \to 0$, which causes the divergence of $\sigma_d$. (b) FI of SLIVER(A)+DD(R) or Polar-SLIVER. SLIVER(A) indicates the sum of FIs from symmetric and antisymmetric channels. The total FI remains non-zero at $d = 0$, which ensures no divergence in $\sigma_d$. We also observe that due to the increase of information in DD(R), the maximum of the total FI occurs at $d \sim 0.7\sigma_{\mathrm{psf}}$. (c) FI of DD(A)+DD(R). We observe that the total FI reduces to zero as $d \to 0$, leading to divergence of $\sigma_d$. The total FI shows a strong local peak at $d \sim 0.9\sigma_{\mathrm{psf}}$. (d) FI of SLIVER+DD for Gaussian PSFs. The FI from SLIVER is a constant over all separations. If all photons were detected by SLIVER, the FI of SLIVER is $F_{\infty,\mathrm{QFI}} = 1/4\sigma^2_{\mathrm{psf}}$, which is the corresponding QFI for Gaussian PSFs.

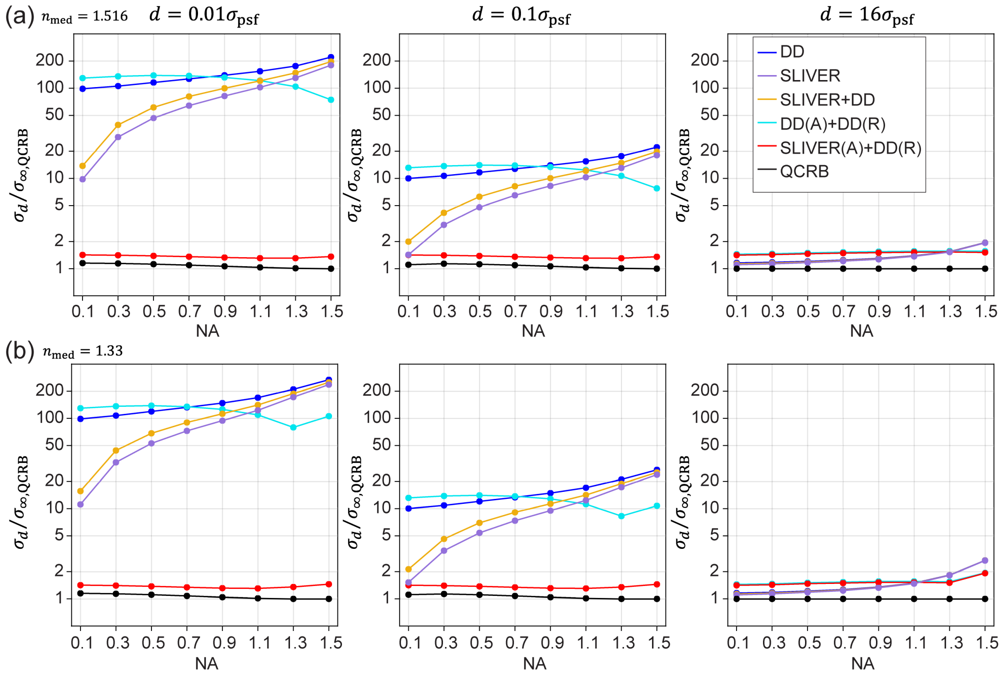


**Fig. S3.** Effect of numerical aperture on separation estimation precision. (a) Comparison of $\sigma_d$ versus numerical aperture for different detection methods when the refractive index is matched. Results at separations of $0.01\sigma_{\text{psf}}$, $0.1\sigma_{\text{psf}}$ and $16\sigma_{\text{psf}}$ are shown. (b) Same as (a), but with index mismatching. Simulation parameters are the same as those defined in Fig. 3(a).

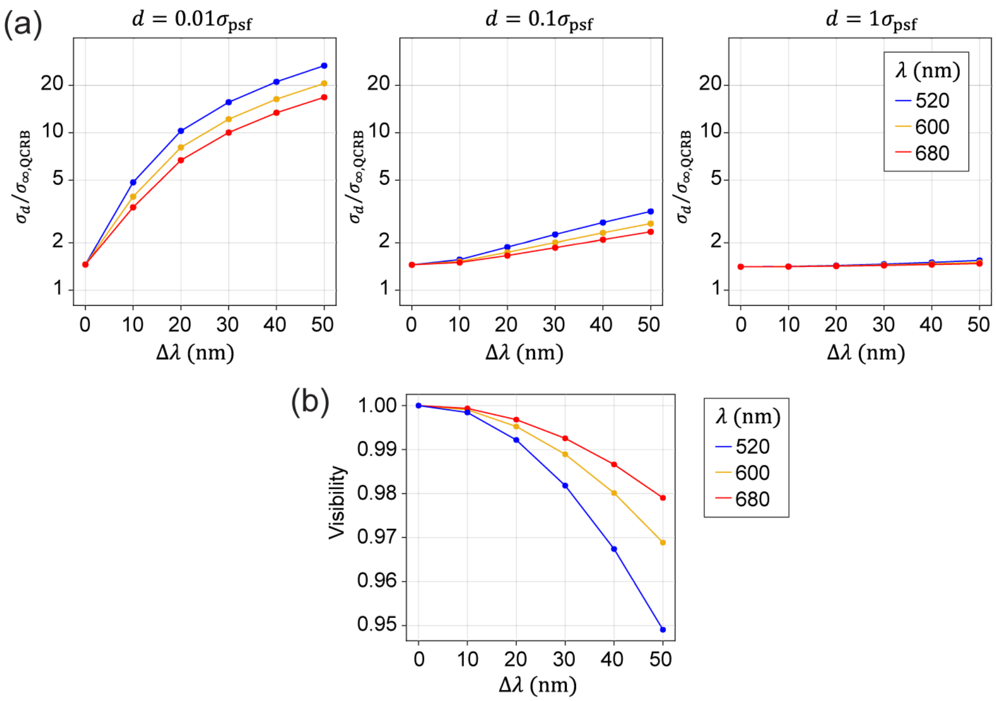


**Fig. S4.** Comparison of separation estimation for Polar-SLIVER with VWP optimized at different central wavelengths of the emission filters. (a) Precision of separation estimation versus detection bandwidth. Results at separations of $0.01\sigma_{\text{psf}}$, $0.1\sigma_{\text{psf}}$ and $1\sigma_{\text{psf}}$ are shown. $\sigma_d$ starts to diverge as $d \to 0$ with non-zero detection bandwidth. The divergence is larger at shorter wavelengths. (b) Visibility versus detection bandwidth. Visibility reduces with increasing detection bandwidth, and the reduction is higher for shorter wavelengths. Simulation parameters are: $n_a = 1.45$, $n_{\text{med}} = 1.33$.

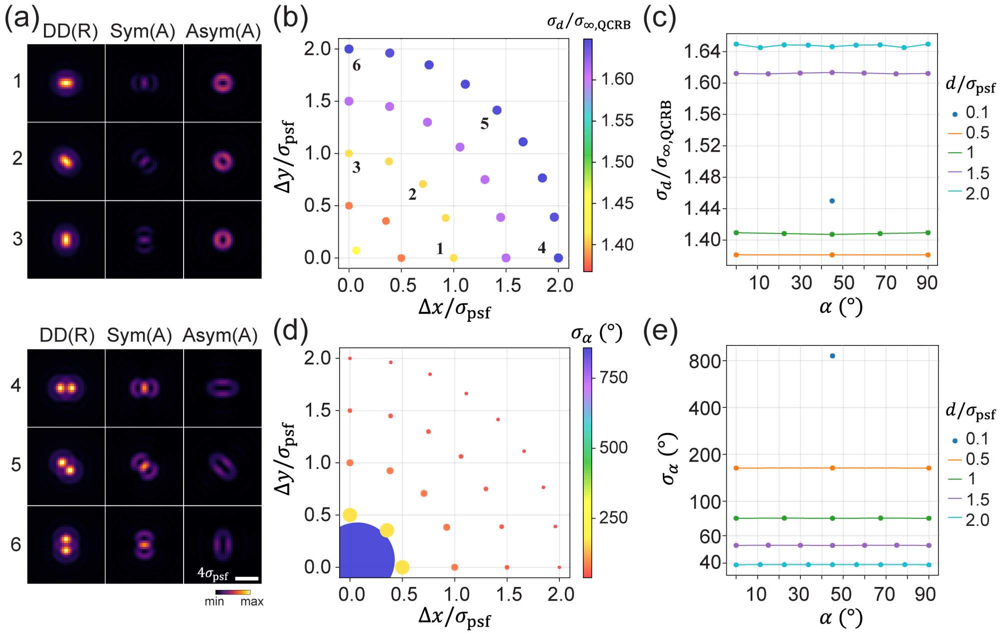


**Fig. S5.** Estimation precisions of angle and separation using Polar-SLIVER. (a) PSFs at separations of $1\sigma_{\text{psf}}$ and $2\sigma_{\text{psf}}$, and three angles of 0°, 45° and 90°. (b) Scatter plot of separation estimation precision $\sigma_d$ over separations along the x and y axes. Labels 1-6 correspond to PSFs in (a). Color and size are scaled with $\sigma_d/\sigma_{\infty,\text{QCRB}}$. (c) $\sigma_d$ versus angle. Each point corresponds to one point in (b). $\sigma_d$ has little dependence on the angle of the two-emitter pair. (d) Scatter plot of angle estimation precision $\sigma_\alpha$ over separations along the x and y axes. Color and size are scaled with $\sigma_\alpha$. (e) $\sigma_\alpha$ versus angle. Each point corresponds to one point in (d). $\sigma_\alpha$ also has little dependence on the angle of the two-emitter pair. However, $\sigma_\alpha$ diverges at small separations. Precisions were obtained under full-parameter estimation. Simulation parameters are: $n_a = 1.45$, $\lambda = 680$ nm, $n_{\text{med}} = 1.33$, VWP is ideal.

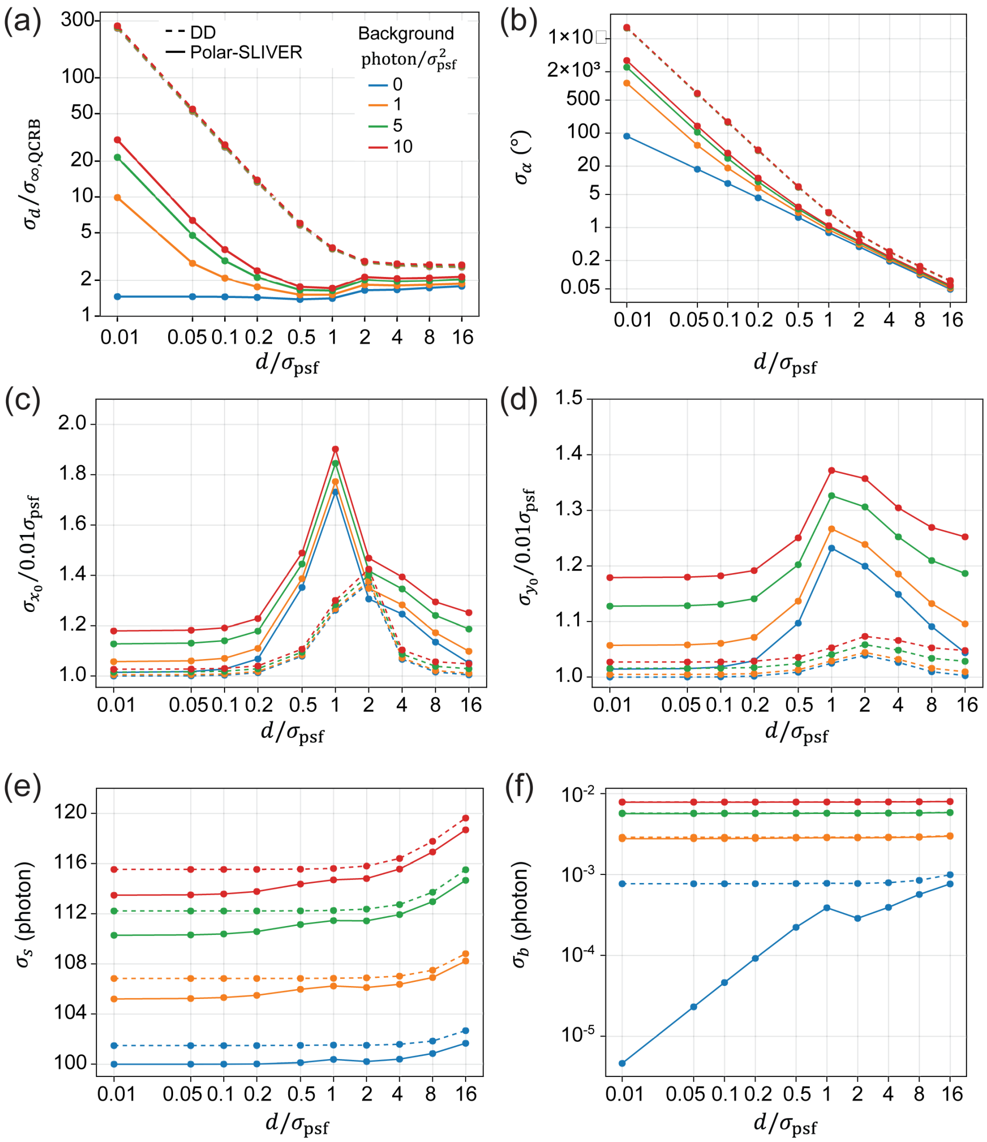


**Fig. S6.** Estimation precision of all parameters for Polar-SLIVER and DD at various background levels. (a-f) Estimation precision for separation, angle, centroid position $(x_0, y_0)$, total photon, and background photon from full parameter estimation. DD is less affected by the background. Background will induce divergence of $\sigma_d$ and increase the divergence of $\sigma_\alpha$ for Polar-SLIVER. Simulation parameters are: $n_a = 1.45$, $\lambda = 680$ nm, $n_{\rm med} = 1.33$, total photon $s = 10000$, VWP is ideal.

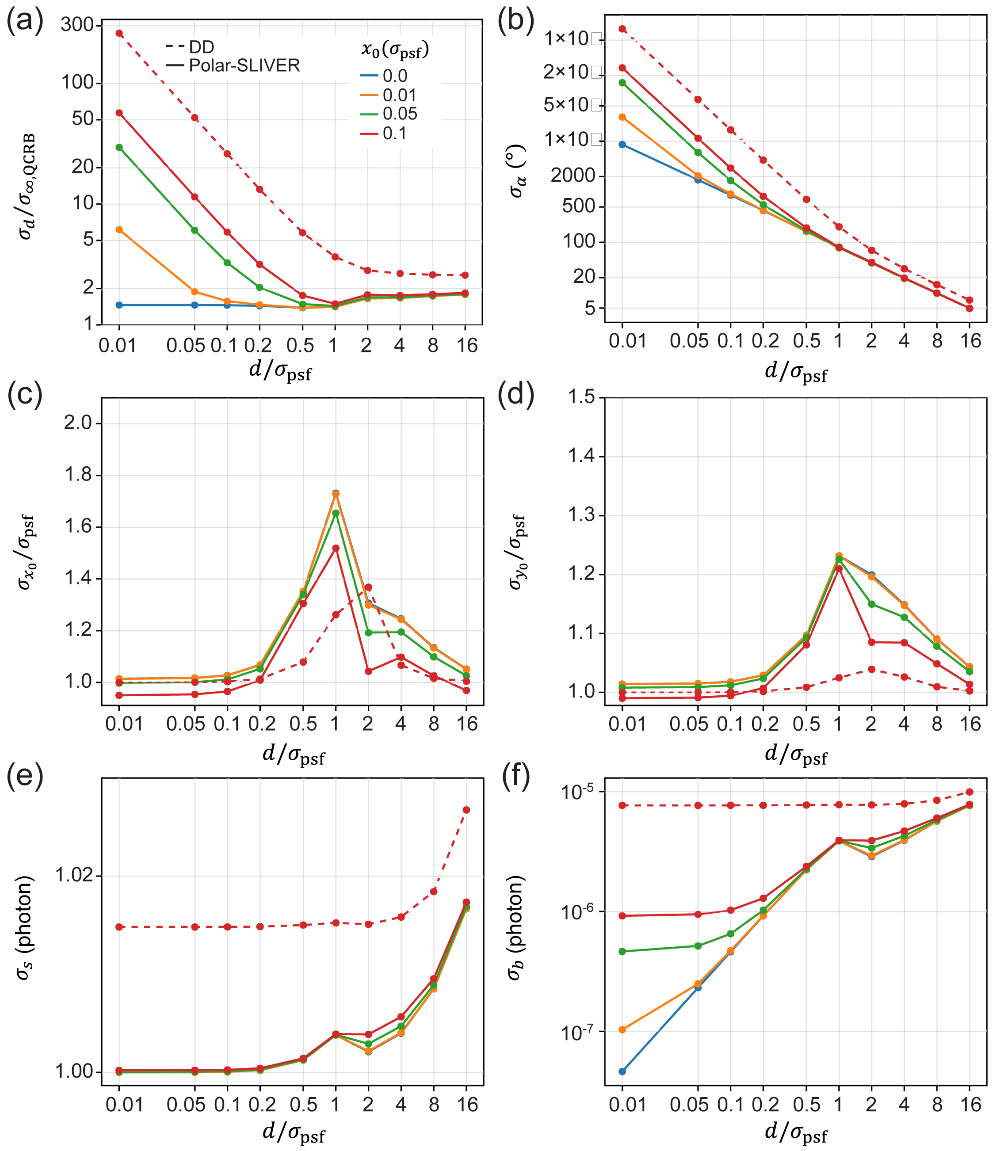


**Fig. S7.** Estimation precision of all parameters for Polar-SLIVER and DD at different amounts of misalignment. Misalignment is introduced by adding an offset to the centroid position along the x-axis ($x_0$). (a-f) Estimation precision for separation, angle, centroid position ($x_0, y_0$), total photon, and background photon from full parameter estimation. As misalignment doesn't apply to DD, misalignment only affects Polar-SLIVER. Misalignment will induce divergence of $\sigma_d$ and increase the divergence of $\sigma_\alpha$ for Polar-SLIVER. Simulation parameters are: $n_a = 1.45$, $\lambda = 680$ nm, $n_{\rm med} = 1.33$, VWP is ideal.

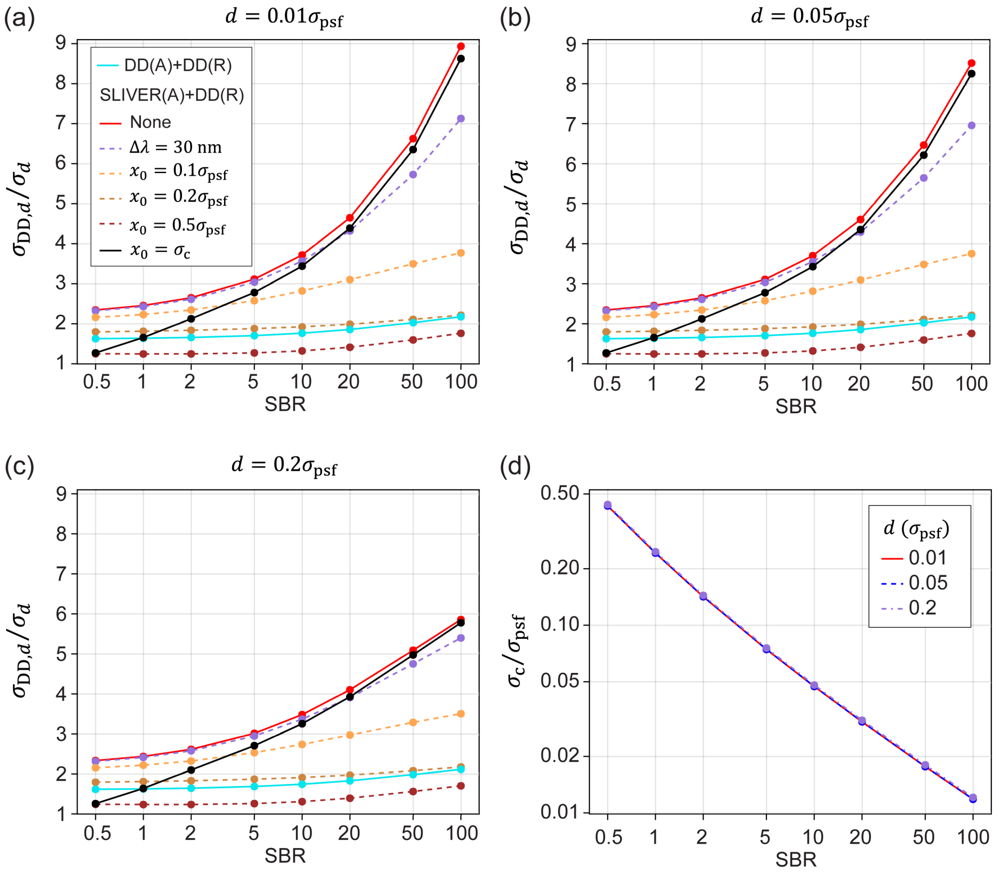


**Fig. S8.** Separation estimation precision at different levels of signal-to-background ratio (SBR). (a-c) Separation estimation precision $\sigma_d$ versus SBR relative to the one achieved by DD, $\sigma_{DD,d}$. The value indicates the magnitude of improvement from DD. Results at separations of $0.01\sigma_{psf}$, $0.05\sigma_{psf}$ , and $0.2\sigma_{psf}$ are shown. With zero detection bandwidth and misalignment, SLIVER(A)+DD(R) or Polar-SLIVER is always >2 times better than DD (red solid line). Misalignment will reduce the performance of Polar-SLIVER to that of DD(A)+DD(R) and DD at a misalignment of $0.2\sigma_{psf}$ and $0.5\sigma_{psf}$, respectively. However, if the misalignment is only due to the precision of the centroid estimation ($\sigma_c$), Polar-SLIVER continues to outperform the other methods when SBR>1 (black solid line). (d) Centroid estimation precision versus SBR using Polar-SLIVER. All three lines overlap, indicating that $\sigma_c$ has minimal dependence on separation.

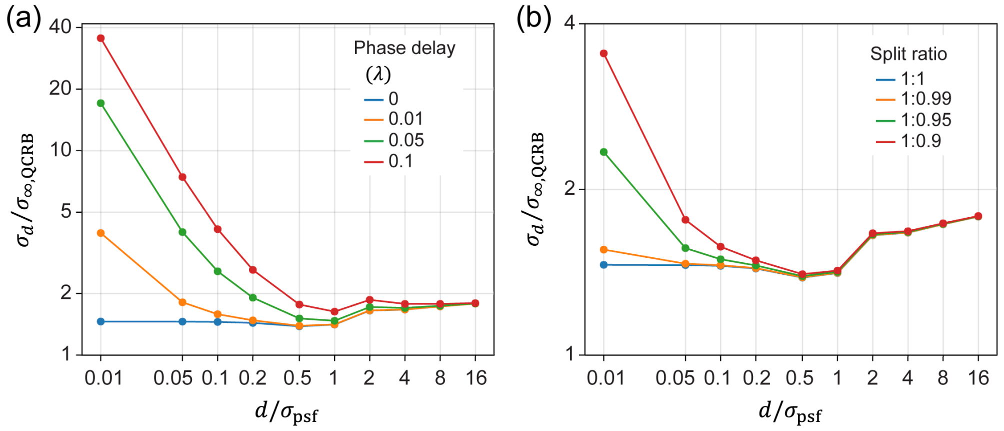


**Fig. S9.** Effect of phase delay and intensity split ratio between two interference arms on separation estimation for Polar-SLIVER. (a) Separation estimation precision $\sigma_d$ versus separation at various phase delays. Phase delay will cause divergence of $\sigma_d$ as $d \to 0$. However, a phase delay of $0.01\lambda$ is readily achievable with existing phase stabilization methods. (b) $\sigma_d$ versus separation at various split ratios. Unequal split ratio will also induce divergence of $\sigma_d$ as $d \to 0$. However, the effect is small. Simulation parameters are: $n_a = 1.45$, $\lambda = 680$ nm, $n_{\text{med}} = 1.33$, VWP is ideal. Precisions were calculated under full parameter estimation.

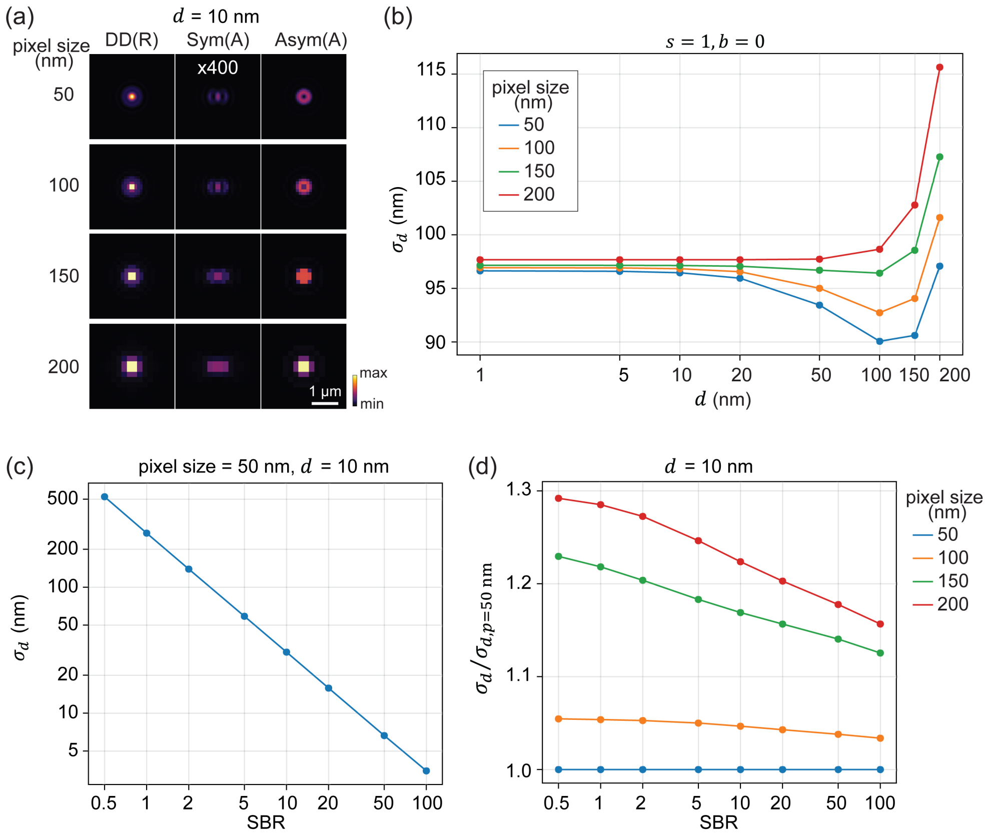


**Fig. S10.** Effect of pixel size on separation estimation using Polar-SLIVER. (a) Emission patterns of two emitters at different pixel sizes. The separation of the two emitters is 10 nm. Here we use the physical unit for both $d$ and $\sigma_d$, because $\sigma_{\mathrm{psf}}$ is pixel size dependent. Each pixel value is calculated as the integral over 2×2, 4×4, 6×6, and 8×8 up-sampling points within the pixel size of 50 nm, 100 nm, 150 nm, and 200 nm, respectively. The image size is kept to around 3.2×3.2 $\mu m^2$ for different pixel sizes. (b) $\sigma_d$ versus $d$ at different pixel sizes. Results were obtained under the ideal condition. $\sigma_d$ has little dependence on pixel size at $d < 20$ nm. At $d > 20$ nm, a smaller pixel size performs better, but the differences are still small. (c) $\sigma_d$ versus SBR at a pixel size of 50 nm and a separation of 10 nm. (d) $\sigma_d$ versus SBR at different pixel sizes and a separation of 10 nm. Precision values are reported relative to those obtained at a pixel size of 50 nm. Larger pixel sizes (150 nm and 200 nm) have lower tolerance to SBR. A pixel size of 100 nm is close to the Nyquist rate and achieves comparable performance with a pixel size of 50 nm. Simulation parameters are: $n_a = 1.45$, $\lambda = 680$ nm, $n_{\mathrm{med}} = 1.33$, VWP is ideal

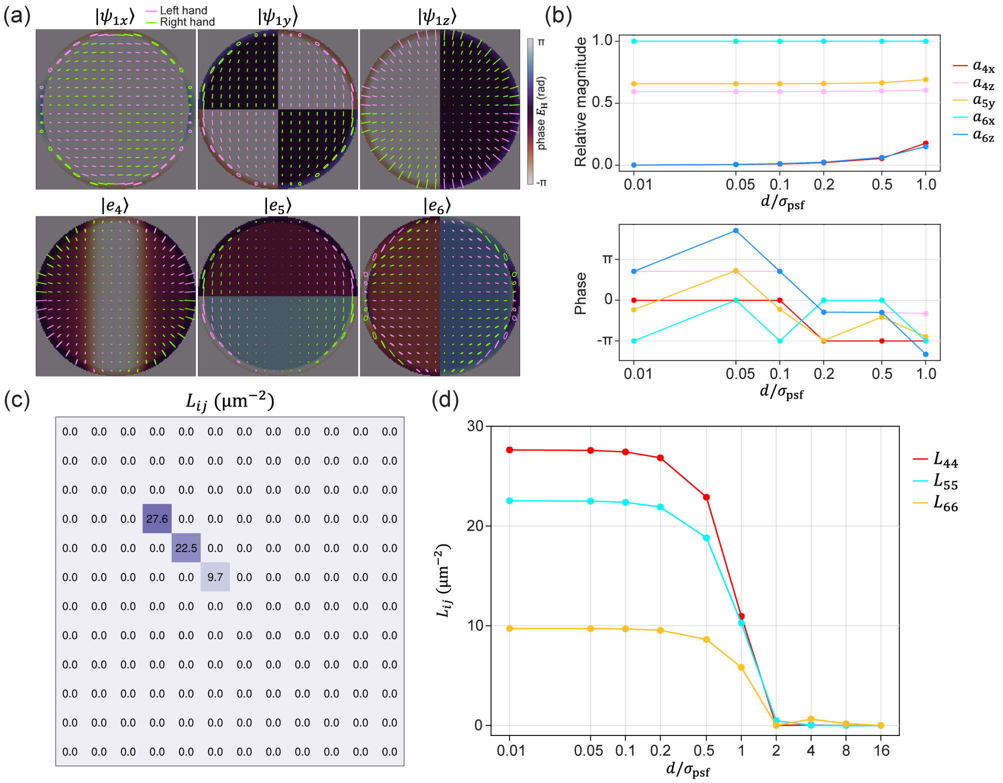


**Fig. S11.** Example QFI calculation using vector fields. (a) Vector representation of the electric fields at the pupil plane from emitter 1 (top row), and the eigenstates 4, 5, 6 (bottom row). The separation of the two dipole emitters is $0.01\sigma_{\text{psf}}$. The polarization state at a certain pupil position is illustrated as right-handed (green) or left-handed (magenta) elliptical polarization. The background heatmap scales with the phase of the horizontal polarization component ($E_{\text{H}}$) of each field. (b) Coefficients of constructing the eigenstates 4, 5, 6 in terms of dipole fields, defined in Eqs. (S133-S135). The magnitudes of the coefficients are shown as the relative values to the magnitude of $a_{6x}$ at each separation. (c) Element-wise QFI matrix $L$ as defined in Eq. (S132). The separation of the two dipole emitters is $0.01\sigma_{\text{psf}}$. (d) Elements $L_{44}, L_{55}, L_{66}$ in matrix $L$ at different separations. Simulation parameters are: $n_a = 1.45$, $\lambda = 680$ nm, $n_{\text{med}} = 1.33$.

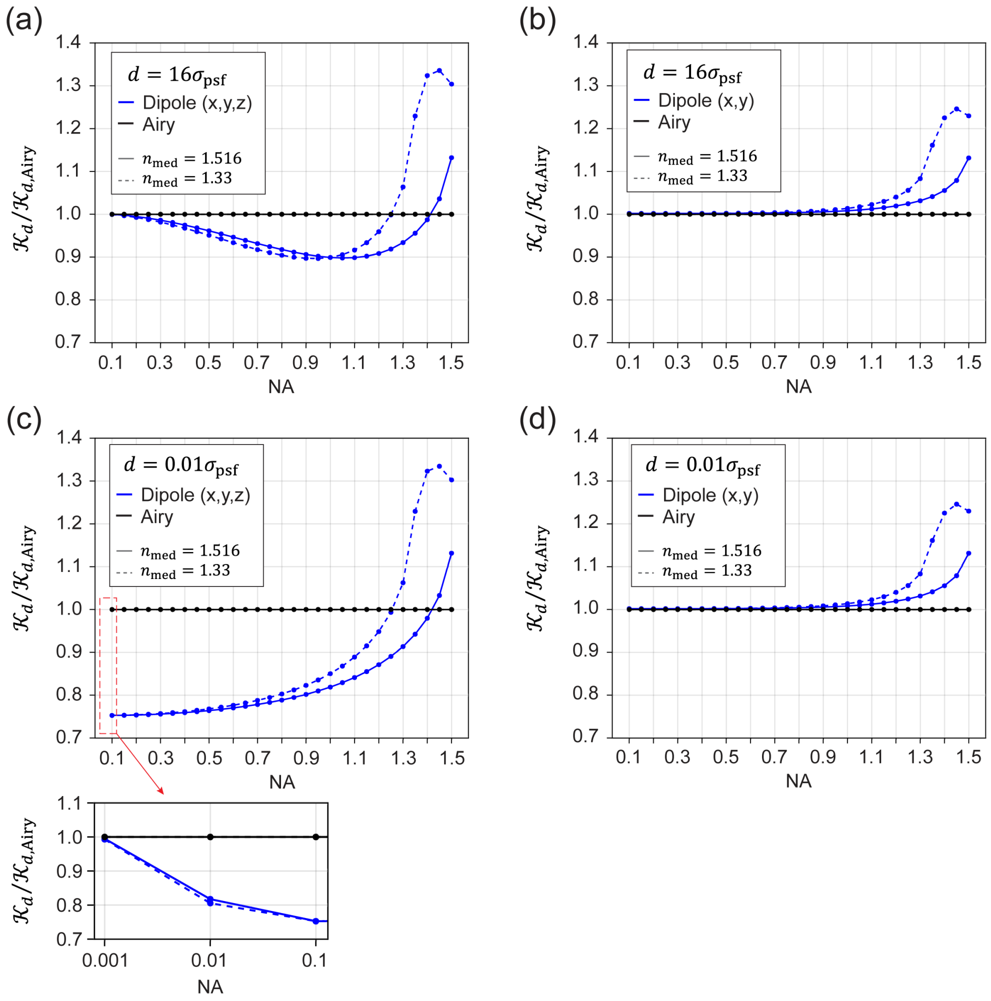


**Fig. S12.** Comparison of QFI for separation estimation of dipole emitters and scalar point sources. (a, c) QFI of freely rotating dipoles with a separation of $16\sigma_{\mathrm{psf}}$ and $0.01\sigma_{\mathrm{psf}}$, respectively. (b, d) QFI of freely rotating dipoles (lateral only) with a separation of $16\sigma_{\mathrm{psf}}$ and $0.01\sigma_{\mathrm{psf}}$, respectively. Here, the dipole orientation is confined in the lateral plane. All QFIs are normalized to the QFI of scalar point sources. For the scalar point source, we define its pupil function as a circular aperture, whose PSF is the Airy pattern. For each condition, results under refractive index-matched ($n_{\mathrm{med}} = 1.516$) and mismatched ($n_{\mathrm{med}} = 1.33$) conditions are shown.

### Supplementary Text

Throughout the text, we used the following convention: bold letter represents a vector, italic letter represents a variable, and regular letter represents a constant or a descriptive subscript.

#### 1. ELECTRIC FIELD FROM DIPOLE RADIATION

Here, we derive the electric field from the dipole radiation of a static dipole. The coordinate system is defined in Fig. 1(d). Using the Weyl identity [1–4], we can express the dipole radiation in its plane wave representation:

$$E_\theta = \boldsymbol{e}_\theta \cdot \boldsymbol{\mu}, \tag{S1}$$

$$E_\varphi = \boldsymbol{e}_\varphi \cdot \boldsymbol{\mu}. \tag{S2}$$

The electric field propagates along the wave vector $\boldsymbol{k} = 2\pi n(\boldsymbol{e}_\theta \times \boldsymbol{e}_\varphi)/\lambda$. The amplitudes of the electric fields, $E_\theta$ and $E_\varphi$, are the electric field components parallel and perpendicular to the incident plane, which is defined by $\boldsymbol{k}$ and the optical axis (z-axis). The dipole moment of a single dipole is defined by $\boldsymbol{\mu} = [\mu_x, \mu_y, \mu_z]$. The vectors $\boldsymbol{e}_\theta = [\cos\theta\cos\varphi, \cos\theta\sin\varphi, -\sin\theta]$ and $\boldsymbol{e}_\varphi = [-\sin\varphi, \cos\varphi, 0]$ are unit vectors in the directions of increasing angles $\theta$ and $\varphi$ in a spherical coordinate, respectively. After transmitting through a multi-layer medium (sample, coverslip and immersion mediums) and the objective lens, the electric field at the back focal plane of the objective is,

$$E_\mathrm{R} = aT_\theta E_\theta, \tag{S3}$$

$$E_\mathrm{A} = aT_\varphi E_\varphi, \tag{S4}$$

where $T_\theta$ and $T_\varphi$ are the Fresnel transmission coefficients [5],

$$T_\theta = \frac{2n_1\cos\theta_1}{n_1\cos\theta_2 + n_2\cos\theta_1}\frac{2n_2\cos\theta_2}{n_2\cos\theta_3 + n_3\cos\theta_2}, \tag{S5}$$

$$T_\varphi = \frac{2n_1\cos\theta_1}{n_1\cos\theta_1 + n_2\cos\theta_2}\frac{2n_2\cos\theta_2}{n_2\cos\theta_2 + n_3\cos\theta_3}, \tag{S6}$$

The subscripts 1, 2 and 3 denote the sample, coverslip and immersion mediums, $n$ denotes the refractive index and $\theta$ is the angle between $\boldsymbol{k}$ and the z-axis. The apodization term $a = \sqrt{\cos\theta_3}/\cos\theta_1$ arises from the conversion of a spherical wavefront to a plane wavefront after transmitting through the objective [6, 7]. We define $E_\mathrm{R}$ and $E_\mathrm{A}$ as the radial and azimuthal components of the dipole radiation from a single dipole oscillating along $\boldsymbol{\mu}$.

#### 2. PUPIL AFTER A VORTEX HALF-WAVE PLATE

The Jones matrix for a vortex half-wave plate (VWP) can be written as [8–11]:

$$\mathcal{M} = \begin{bmatrix} \cos\frac{\delta}{2} + i\sin\frac{\delta}{2}\cos\varphi & i\sin\frac{\delta}{2}\sin\varphi \\ i\sin\frac{\delta}{2}\sin\varphi & \cos\frac{\delta}{2} - i\sin\frac{\delta}{2}\cos\varphi \end{bmatrix}, \tag{S7}$$

where $\delta = 2\pi(n_e - n_o)t/\lambda$, with $t$ being the thickness of the VWP. The Jones vectors for the radial and azimuthal polarizations are [9]:

$$\mathbf{R} = \begin{bmatrix} \cos\varphi \\ \sin\varphi \end{bmatrix}, \mathbf{A} = \begin{bmatrix} -\sin\varphi \\ \cos\varphi \end{bmatrix}. \tag{S8}$$

And the Jones vectors for the horizontal and vertical polarizations are [9]:

$$\mathbf{H} = \begin{bmatrix} 1 \\ 0 \end{bmatrix}, \mathbf{V} = \begin{bmatrix} 0 \\ 1 \end{bmatrix}. \tag{S9}$$

The radially and azimuthally polarized fields after the VWP become

$$\boldsymbol{E}_{\text{VWP,R}} = E_{\text{R}}\mathcal{M}\mathbf{R} = E_{\text{R}} \cos\frac{\delta}{2}\mathbf{R} + iE_{\text{R}} \sin\frac{\delta}{2}\mathbf{H}, \tag{S10}$$

$$\boldsymbol{E}_{\text{VWP,A}} = E_{\text{A}}\mathcal{M}\mathbf{A} = E_{\text{A}} \cos\frac{\delta}{2}\mathbf{A} - iE_{\text{A}} \sin\frac{\delta}{2}\mathbf{V}. \tag{S11}$$

Through a polarized beam splitter (PBS), the above fields can be projected onto horizontal and vertical polarizations,

$$h_{\text{VWP,H}} = (\boldsymbol{E}_{\text{VWP,R}} + \boldsymbol{E}_{\text{VWP,A}}) \cdot \mathbf{H} = (E_{\text{R}} \cos\varphi - E_{\text{A}} \sin\varphi) \cos\frac{\delta}{2} + iE_{\text{R}} \sin\frac{\delta}{2}, \tag{S12}$$

$$h_{\text{VWP,V}} = (\boldsymbol{E}_{\text{VWP,R}} + \boldsymbol{E}_{\text{VWP,A}}) \cdot \mathbf{V} = (E_{\text{R}} \sin\varphi + E_{\text{A}} \cos\varphi) \cos\frac{\delta}{2} - iE_{\text{A}} \sin\frac{\delta}{2}. \tag{S13}$$

We use $h_{\text{VWP,H}}$ and $h_{\text{VWP,V}}$ to represent the pupil functions after the PBS. For SLIVER(A)+DD(R) or Polar-SLIVER, the field $h_{\text{VWP,H}}$ will enter the DD path, while the field $h_{\text{VWP,V}}$ will enter the SLIVER(A) path. The phase delay $\delta$ is a function of the emission wavelength $\lambda$. Assuming the VWP is designed for the central wavelength $\lambda_0$ of an emission band-pass filter, we can express $\delta = \pi + \Delta\delta$, with $\Delta\delta$ being the residue phase delay when $\lambda$ deviates from $\lambda_0$. Then we can rewrite Eqs. (S12, S13) as,

$$h_{\text{VWP,H}} = -(E_{\text{R}} \cos\varphi - E_{\text{A}} \sin\varphi) \sin\frac{\Delta\delta}{2} + iE_{\text{R}} \cos\frac{\Delta\delta}{2}, \tag{S14}$$

$$h_{\text{VWP,V}} = -(E_{\text{R}} \sin\varphi + E_{\text{A}} \cos\varphi) \sin\frac{\Delta\delta}{2} - iE_{\text{A}} \cos\frac{\Delta\delta}{2}. \tag{S15}$$

When $\Delta\delta = 0$, the radial and azimuthal components of the dipole radiation will be directly mapped to horizontally and vertically polarized fields after the PBS, given by $h_{\text{VWP,H}} = iE_{\text{R}}$ and $h_{\text{VWP,V}} = -iE_{\text{A}}$. However, this requires a band-pass filter with zero bandwidth, which is not feasible for fluorescence imaging. In practice, $h_{\text{VWP,H}}$ and $h_{\text{VWP,V}}$ will not originate solely from the radial or azimuthal component of the dipole radiation. On the other hand, if we remove the VWP, equivalent to $\delta = 0$, Eqs. (S12, S13) become,

$$h_{\text{H}} = E_{\text{R}} \cos\varphi - E_{\text{A}} \sin\varphi, \tag{S16}$$

$$h_{\text{V}} = E_{\text{R}} \sin\varphi + E_{\text{A}} \cos\varphi. \tag{S17}$$

We use $h_{\text{H}}$ and $h_{\text{V}}$ to represent the pupil functions when imaging a dipole emitter using the detection methods without a VWP. The above equations show that $h_{\text{H}}$ and $h_{\text{V}}$ comprise both radial and azimuthal components of the dipole radiation, making them suboptimal for SLIVER detection.

### 3. CALCULATION OF THE PSF OF A SINGLE DIPOLE AS A 1D INTEGRAL

The point spread function from each electric field component (e.g. $h_{\text{H}}$ and $h_{\text{V}}$) can be calculated via the Fourier transform of the electric field at the pupil plane [12]:

$$U(x,y) = \mathcal{F}(h) = \iint h(k_x,k_y) e^{2\pi i(k_x x + k_y y)} dk_x dk_y. \tag{S18}$$

where $\mathcal{F}$ denotes a 2D Fourier transform. Convert Eq. (S18) to polar coordinates,

$$U(r,\varphi_{\text{s}}) = \iint h(k_r,\varphi) e^{2\pi i k_r r \cos(\varphi - \varphi_{\text{s}})} k_r dk_r d\varphi, \tag{S19}$$

where $r = \sqrt{x^2 + y^2}$ , $k_r = \sqrt{k_x^2 + k_y^2}$. Here, $\varphi_{\text{s}}$ denotes the polar angle at the image plane. We observe that the electric fields in Eqs. (S14-S17) can be written in the form of $h = \sum_i R_i(\theta) A_i(\varphi)$, where $i$ indicates the term index. The terms $A_i(\varphi)$ are functions of $\cos\varphi$ and $\sin\varphi$, whose integral over $\varphi$ can be expressed in terms of Bessel functions [12]. Here we define the operation $\mathcal{A}(h)$ as a 1D integral over $\varphi$:

$$\mathcal{A}(h) = \int_0^{2\pi} h e^{2\pi i k_r r \cos(\varphi - \varphi_s)} d\varphi. \tag{S20}$$

The operations of $\mathcal{A}$ to each possible $A_i(\varphi)$ are listed below:

$$I_0 = \mathcal{A}(1) = 2\pi J_0(2\pi k_r r), \tag{S21}$$
$$I_1 = \mathcal{A}(\sin\varphi) = 2\pi i J_1(2\pi k_r r)\sin\varphi_s, \tag{S22}$$
$$I_2 = \mathcal{A}(\cos\varphi) = 2\pi i J_1(2\pi k_r r)\cos\varphi_s, \tag{S23}$$
$$I_3 = \mathcal{A}(\cos^2\varphi) = \pi[J_0(2\pi k_r r) - J_2(2\pi k_r r)\cos 2\varphi_s], \tag{S24}$$
$$I_4 = \mathcal{A}(\sin^2\varphi) = \pi[J_0(2\pi k_r r) + J_2(2\pi k_r r)\cos 2\varphi_s], \tag{S25}$$
$$I_5 = \mathcal{A}(\sin\varphi\cos\varphi) = -\pi J_2(2\pi k_r r)\sin 2\varphi_s. \tag{S26}$$

We use $J_m$ to indicate the Bessel function of the first kind. Given the above integrals, we can calculate the Fourier transform of the electric fields $h_\mathrm{H}$, $h_\mathrm{V}$, $E_\mathrm{R}$ and $E_\mathrm{A}$ as a 1D integral:

$$\tilde{h}_\mathrm{H} = \mathcal{F}(h_\mathrm{H}) = \int_0^{n_a/\lambda} \left(aT_\theta[I_3\cos\theta, I_5\cos\theta, -I_2\sin\theta]\cdot\boldsymbol{\mu} + aT_\varphi[I_4, -I_5, 0]\cdot\boldsymbol{\mu}\right) k_r dk_r, \tag{S27}$$
$$\tilde{h}_\mathrm{V} = \mathcal{F}(h_\mathrm{V}) = \int_0^{n_a/\lambda} \left(aT_\theta[I_5\cos\theta, I_4\cos\theta, -I_1\sin\theta]\cdot\boldsymbol{\mu} + aT_\varphi[-I_5, I_3, 0]\cdot\boldsymbol{\mu}\right) k_r dk_r, \tag{S28}$$
$$\tilde{E}_\mathrm{R} = \mathcal{F}(E_\mathrm{R}) = \int_0^{n_a/\lambda} aT_\theta[I_2\cos\theta, I_1\cos\theta, -I_0\sin\theta]\cdot\boldsymbol{\mu} k_r dk_r, \tag{S29}$$
$$\tilde{E}_\mathrm{A} = \mathcal{F}(E_\mathrm{A}) = \int_0^{n_a/\lambda} aT_\varphi[-I_1, I_2, 0]\cdot\boldsymbol{\mu} k_r dk_r, \tag{S30}$$

where $\sin\theta = k_r\lambda/n$ and $\cos\theta = \sqrt{1-(k_r\lambda/n)^2}$ are functions of $k_r$, with $n$ being the refractive index of the sample medium. Then the Fourier transform of the electric fields at the pupil plane for detection methods with a VWP, $h_\mathrm{VWP,H}$ and $h_\mathrm{VWP,V}$, can be written as:

$$\tilde{h}_\mathrm{VWP,H} = \mathcal{F}(h_\mathrm{VWP,H}) = -\tilde{h}_\mathrm{H}\sin\frac{\Delta\delta}{2} + i\tilde{E}_\mathrm{R}\cos\frac{\Delta\delta}{2}, \tag{S31}$$
$$\tilde{h}_\mathrm{VWP,V} = \mathcal{F}(h_\mathrm{VWP,V}) = -\tilde{h}_\mathrm{V}\sin\frac{\Delta\delta}{2} - i\tilde{E}_\mathrm{A}\cos\frac{\Delta\delta}{2}. \tag{S32}$$

### 4. CALCULATION OF THE PSF OF A FREELY ROTATING DIPOLE

A freely rotating dipole is defined as one that can freely rotate to any orientation within the fluorescent lifetime of a single fluorophore. Since the dipole can orient in any direction before emitting a photon, the emitted photon will have no memory of the excitation polarization. Therefore, the excitation field can be considered as a scalar field, and the dipole moment is equal at all orientations. Given that the camera integration time is much longer than the fluorescent lifetime, the PSF of a freely rotating dipole will be the incoherent sum of contributions from all possible dipole orientations [13]. Given the electric field at the pupil plane, for example $h_\mathrm{H}$, the PSF of a freely rotating dipole can be calculated from,

$$I = \sum_s |\mathcal{F}(h_{\mathrm{H},s})|^2, \tag{S33}$$

where $s$ indexes all dipole orientations and $I$ denotes the intensity PSF. From Eq. (S16), we can rewrite $h_\mathrm{H}$ as,

$$h_\mathrm{H} = \sum_j \omega_j \mu_{js}, \quad j = \mathrm{x}, \mathrm{y}, \mathrm{z} \tag{S34}$$

where $\omega_j = aT_\theta e_{\theta j}\cos\varphi - aT_\varphi e_{\varphi j}\sin\varphi$, and $\mu_{js}$ denotes the Cartesian components of the dipole moment at orientation $s$. Then we can rewrite Eq. (S33) as,

$$I = \sum_s \left|\mathcal{F}\left(\sum_j \omega_j\mu_{js}\right)\right|^2 = \sum_s \left|\sum_j \mathcal{F}\left(\omega_j\right)\mu_{js}\right|^2. \tag{S35}$$

We define $\tilde{\omega}_j = \mathcal{F}(\omega_j)$. By explicitly calculating the modulus square, we have

$$\sum_s 2\mathrm{Re}(\tilde{\omega}_m\tilde{\omega}_n^*)\mu_{ms}\mu_{ns} = 0, \quad m \neq n. \tag{S36}$$

The above equation is true only when the dipole moment is equal at all orientations and $s$ uniformly samples all dipole orientations. Then Eq. (S35) can be simplified to

$$I = \sum_s \sum_j |\tilde{\omega}_j|^2 \mu_{js}^2 = \sum_j |\tilde{\omega}_j|^2 \sum_s \mu_{js}^2. \tag{S37}$$

For isotropic dipole orientations, we have $\sum_s \mu_{js}^2 = C/3$, where $C$ is a constant. Then we have

$$I = C \sum_j |\tilde{\omega}_j|^2, \quad j = \mathrm{x, y, z}. \tag{S38}$$

The above equation indicates that the PSF of a freely rotating dipole is equivalent to the sum of the PSFs from three dipoles oriented along x, y, and z directions.

### 5. CRB OF POLAR-SLIVER

#### A. Symmetric channel

Based on the definition of the two-emitter model in the main text, the two-emitter model of SLIVER(A) in the symmetric channel can be written as:

$$I_{\mathrm{2em,sym,A}} = \sum_{j=\mathrm{x,y}} \frac{1}{4} \left| \tilde{\omega}_{j\mathrm{A}} \left( x - \frac{d}{2} \right) + \tilde{\omega}_{j\mathrm{A}} \left( -x - \frac{d}{2} \right) \right|^2 + \frac{1}{4} \left| \tilde{\omega}_{j\mathrm{A}} \left( x + \frac{d}{2} \right) + \tilde{\omega}_{j\mathrm{A}} \left( -x + \frac{d}{2} \right) \right|^2 \tag{S39}$$

Here $j = \mathrm{x, y}$ denotes the dipole oriented along the x and y directions. We use a regular font for x and y to distinguish them from the variable $x$ and $y$. Since $\tilde{\omega}_{j\mathrm{A}}$ are odd functions, we can rewrite the above equation as,

$$I_{\mathrm{2em,sym,A}} = \sum_{j=\mathrm{x,y}} \frac{1}{2} \left| \tilde{\omega}_{j\mathrm{A}} \left( x - \frac{d}{2} \right) - \tilde{\omega}_{j\mathrm{A}} \left( x + \frac{d}{2} \right) \right|^2 = \sum_{j=\mathrm{x,y}} \frac{1}{2} \left| \tilde{\omega}_{j\mathrm{A1}} - \tilde{\omega}_{j\mathrm{A2}} \right|^2 \tag{S40}$$

We introduce $\tilde{\omega}_{j\mathrm{A1}}$ and $\tilde{\omega}_{j\mathrm{A2}}$ as defined above to simplify the following derivations. For now, we consider $\tilde{\omega}_{j\mathrm{A}}$ to be real functions, which is true when the refractive indices of the sample and immersion mediums are equal. Then we can rewrite Eq. (S40) as

$$I_{\mathrm{2em,sym,A}} = \sum_{j=\mathrm{x,y}} \frac{1}{2} \left( \tilde{\omega}_{j\mathrm{A1}} - \tilde{\omega}_{j\mathrm{A2}} \right)^2. \tag{S41}$$

The derivatives of $I_{\mathrm{2em,sym,A}}$ with respect to $d$ is,

$$\frac{\partial I_{\mathrm{2em,sym,A}}}{\partial d} = \sum_{j=\mathrm{x,y}} \frac{1}{2} (\tilde{\omega}_{j\mathrm{A2}} - \tilde{\omega}_{j\mathrm{A1}}) \left( \frac{\partial \tilde{\omega}_{j\mathrm{A1}}}{\partial x} + \frac{\partial \tilde{\omega}_{j\mathrm{A2}}}{\partial x} \right) \tag{S42}$$

The Fisher information for estimating $d$ is given by,

$$F_d = \iint \frac{1}{I_{\mathrm{2em,sym,A}}} \left( \frac{\partial I_{\mathrm{2em,sym,A}}}{\partial d} \right)^2 dx dy. \tag{S43}$$

We define the integrand as,

$$f_{\mathrm{sym},d} = \frac{1}{I_{\mathrm{2em,sym,A}}} \left( \frac{\partial I_{\mathrm{2em,sym,A}}}{\partial d} \right)^2. \tag{S44}$$

According to Eq. (S40), we have $\tilde{\omega}_{j\mathrm{A1}} \approx \tilde{\omega}_{j\mathrm{A2}}$ as $d \to 0$. In this case, both the denominator and the numerator of $f_{\mathrm{sym},d}$ approach zero. To calculate the limit at $d \to 0$, we apply the L'Hôpital's rule,

$$\lim_{d \to 0} f_{\mathrm{sym},d} = \lim_{d \to 0} \frac{2 \frac{\partial I_{\mathrm{2em,sym,A}}}{\partial d} \frac{\partial^2 I_{\mathrm{2em,sym,A}}}{\partial d^2}}{\frac{\partial I_{\mathrm{2em,sym,A}}}{\partial d}} = \lim_{d \to 0} 2 \frac{\partial^2 I_{\mathrm{2em,sym,A}}}{\partial d^2}. \tag{S45}$$

From Eq. (S42), we have

$$\frac{\partial^2 I_{\mathrm{2em,sym,A}}}{\partial d^2} = \sum_{j=\mathrm{x,y}} \frac{1}{4} \left( \frac{\partial \tilde{\omega}_{j\mathrm{A1}}}{\partial x} + \frac{\partial \tilde{\omega}_{j\mathrm{A2}}}{\partial x} \right)^2 = \sum_{j=\mathrm{x,y}} \left( \frac{\partial \tilde{\omega}_{j\mathrm{A}}}{\partial x} \right)^2. \tag{S46}$$

Insert Eq. (S46) into Eq. (S45), we have

$$\lim_{d\to 0} f_{\text{sym},d} = \sum_{j=\text{x,y}} 2\left(\frac{\partial\tilde{\omega}_{j\text{A}}}{\partial x}\right)^2. \tag{S47}$$

We next consider the limit at $d \to \infty$. The amplitude distribution of the fields $\tilde{\omega}_{j\text{A}}$ is primarily concentrated within a circular area with a radius of $2\sigma_{\text{psf}}$. Therefore, at $d \to \infty$, $\tilde{\omega}_{j\text{A}1}$ and $\tilde{\omega}_{j\text{A}2}$ are well separated spatially, allowing us to approximate $\tilde{\omega}_{j\text{A}1}\tilde{\omega}_{j\text{A}2} \approx 0$. Then Eq. (S41) can be written as,

$$I_{\text{2em,sym,A}} = \sum_{j=\text{x,y}} \frac{1}{2}\left(\tilde{\omega}_{j\text{A}1}^2 + \tilde{\omega}_{j\text{A}2}^2\right). \tag{S48}$$

And the derivative of $I_{\text{2em,sym,A}}$ with respect to $d$ is

$$\frac{\partial I_{\text{2em,sym,A}}}{\partial d} = \sum_{j=\text{x,y}} \frac{1}{2}\tilde{\omega}_{j\text{A}2}\frac{\partial\tilde{\omega}_{j\text{A}2}}{\partial x} - \sum_{j=\text{x,y}} \frac{1}{2}\tilde{\omega}_{j\text{A}1}\frac{\partial\tilde{\omega}_{j\text{A}1}}{\partial x}. \tag{S49}$$

Insert Eqs. (S48, S49) into Eq. (S43), and noting that the product of the first and second terms in Eq. (S49) is also close to zero, we obtain

$$f_{\text{sym},d} = \frac{\frac{1}{2}\left(\sum_{j=\text{x,y}} \tilde{\omega}_{j\text{A}1}\frac{\partial\tilde{\omega}_{j\text{A}1}}{\partial x}\right)^2 + \frac{1}{2}\left(\sum_{j=\text{x,y}} \tilde{\omega}_{j\text{A}2}\frac{\partial\tilde{\omega}_{j\text{A}2}}{\partial x}\right)^2}{\sum_{j=\text{x,y}} \tilde{\omega}_{j\text{A}1}^2 + \sum_{j=\text{x,y}} \tilde{\omega}_{j\text{A}2}^2}. \tag{S50}$$

Since the fields with subscripts 1 and 2 are well separated spatially as $d \to \infty$, we can calculate their integration over the spatial dimension independently,

$$F_d = \iint f_{\text{sym},d}\,dxdy = \iint \frac{\frac{1}{2}\left(\sum_{j=\text{x,y}} \tilde{\omega}_{j\text{A}1}\frac{\partial\tilde{\omega}_{j\text{A}1}}{\partial x}\right)^2}{\sum_{j=\text{x,y}} \tilde{\omega}_{j\text{A}1}^2}dxdy + \iint \frac{\frac{1}{2}\left(\sum_{j=\text{x,y}} \tilde{\omega}_{j\text{A}2}\frac{\partial\tilde{\omega}_{j\text{A}2}}{\partial x}\right)^2}{\sum_{j=\text{x,y}} \tilde{\omega}_{j\text{A}2}^2}dxdy. \tag{S51}$$

Observing that the fields with subscripts 1 and 2 differ only by a translation of $d$, the two integration terms in Eq. (S51) are equal. Therefore, we can rewrite Eq. (S51) as,

$$F_d = \iint \frac{\left(\sum_{j=\text{x,y}} \tilde{\omega}_{j\text{A}}\frac{\partial\tilde{\omega}_{j\text{A}}}{\partial x}\right)^2}{\sum_{j=\text{x,y}} \tilde{\omega}_{j\text{A}}^2}dxdy. \tag{S52}$$

Then we obtain $f_{\text{sym},d}$ at $d \to \infty$,

$$\lim_{d\to\infty} f_{\text{sym},d} = \frac{\left(\sum_{j=\text{x,y}} \tilde{\omega}_{j\text{A}}\frac{\partial\tilde{\omega}_{j\text{A}}}{\partial x}\right)^2}{\sum_{j=\text{x,y}} \tilde{\omega}_{j\text{A}}^2}. \tag{S53}$$

From the above equation, we recognize that the information for estimating $d$ at $d \to \infty$ is exactly one-fourth of the information for estimating the centroid position of a single emitter using direct detection of the azimuthal component, DD(A).

#### B. Antisymmetric channel

Similar to Eqs. (S39-S41), we obtain the two-emitter model in the antisymmetric channel,

$$I_{\text{2em,asym,A}} = \sum_{j=\text{x,y}} \frac{1}{2}\left(\tilde{\omega}_{j\text{A}1} + \tilde{\omega}_{j\text{A}2}\right)^2. \tag{S54}$$

The derivative of $I_{\text{2em,asym,A}}$ with respect to $d$ is

$$\frac{\partial I_{\text{2em,asym,A}}}{\partial d} = \sum_{j=\text{x,y}} \frac{1}{2}(\tilde{\omega}_{j\text{A}1} + \tilde{\omega}_{j\text{A}2})\left(\frac{\partial\tilde{\omega}_{j\text{A}2}}{\partial x} - \frac{\partial\tilde{\omega}_{j\text{A}1}}{\partial x}\right). \tag{S55}$$

We define

$$f_{\text{asym},d} = \frac{1}{I_{\text{2em,asym,A}}}\left(\frac{\partial I_{\text{2em,asym,A}}}{\partial d}\right)^2. \tag{S56}$$

When $d \to 0$, the numerator of $f_{\text{asym},d}$ approaches zero, while the denominator,

$$\lim_{d\to 0} I_{\text{2em,asym,A}} = \sum_{j=\text{x,y}} 2\tilde{\omega}_{j\text{A}}^2, \tag{S57}$$

which is non-zero. Therefore, we have

$$\lim_{d\to 0} f_{\text{asym},d} = 0. \tag{S58}$$

In the case of $d \to \infty$, we have

$$I_{\text{2em,asym,A}} = \sum_{j=\text{x,y}} \frac{1}{2}\left(\tilde{\omega}_{j\text{A1}}^2 + \tilde{\omega}_{j\text{A2}}^2\right), \tag{S59}$$

which is equal to that of the symmetric channel as shown in Eq. (S48). Therefore, we have

$$\lim_{d\to\infty} f_{\text{asym},d} = \lim_{d\to\infty} f_{\text{sym},d}. \tag{S60}$$

#### C. Direct detection channel

The two-emitter model in the DD(R) channel can be written as

$$I_{\text{2em,R}} = \sum_{j=\text{x,y,z}} \frac{1}{2}\left|\tilde{\omega}_{j\text{R}}\left(x - \frac{d}{2}\right)\right|^2 + \frac{1}{2}\left|\tilde{\omega}_{j\text{R}}\left(x + \frac{d}{2}\right)\right|^2 = \sum_{j=\text{x,y,z}} \frac{1}{2}\left|\tilde{\omega}_{j\text{R1}}\right|^2 + \frac{1}{2}\left|\tilde{\omega}_{j\text{R2}}\right|^2. \tag{S61}$$

Assuming $\tilde{\omega}_{j\text{R}}$ are real functions, we can rewrite the above equation as

$$I_{\text{2em,R}} = \sum_{j=\text{x,y,z}} \frac{1}{2}\left(\tilde{\omega}_{j\text{R1}}^2 + \tilde{\omega}_{j\text{R2}}^2\right). \tag{S62}$$

The derivative of $I_{\text{2em,R}}$ with respect to $d$ is

$$\frac{\partial I_{\text{2em,R}}}{\partial d} = \sum_{j=\text{x,y,z}} \frac{1}{2}\left(\tilde{\omega}_{j\text{R2}}\frac{\partial \tilde{\omega}_{j\text{R2}}}{\partial x} - \tilde{\omega}_{j\text{R1}}\frac{\partial \tilde{\omega}_{j\text{R1}}}{\partial x}\right). \tag{S63}$$

We define

$$f_{\text{R},d} = \frac{1}{I_{\text{2em,R}}}\left(\frac{\partial I_{\text{2em,R}}}{\partial d}\right)^2. \tag{S64}$$

When $d \to 0$, the numerator of $f_{\text{R},d}$ approaches zero, while the denominator,

$$\lim_{d\to 0} I_{\text{2em,R}} = \sum_{j=\text{x,y,z}} \tilde{\omega}_{j\text{R}}^2, \tag{S65}$$

which is non-zero. Therefore, we have

$$\lim_{d\to 0} f_{\text{R},d} = 0. \tag{S66}$$

In the case of $d \to \infty$, we have

$$f_{\text{R},d} = \frac{\frac{1}{2}\left(\sum_{j=\text{x,y,z}} \tilde{\omega}_{j\text{R1}}\frac{\partial \tilde{\omega}_{j\text{R1}}}{\partial x}\right)^2 + \frac{1}{2}\left(\sum_{j=\text{x,y,z}} \tilde{\omega}_{j\text{R2}}\frac{\partial \tilde{\omega}_{j\text{R2}}}{\partial x}\right)^2}{\sum_{j=\text{x,y,z}} \tilde{\omega}_{j\text{R1}}^2 + \sum_{j=\text{x,y,z}} \tilde{\omega}_{j\text{R2}}^2}. \tag{S67}$$

Similar to Eqs. (S50-S53), we obtain

$$\lim_{d\to\infty} f_{\text{R},d} = \frac{\left(\sum_{j=\text{x,y,z}} \tilde{\omega}_{j\text{R}}\frac{\partial \tilde{\omega}_{j\text{R}}}{\partial x}\right)^2}{\sum_{j=\text{x,y,z}} \tilde{\omega}_{j\text{R}}^2}. \tag{S68}$$

From the above equation, we recognize that the information for estimating $d$ at $d \to \infty$ is exactly one-fourth of the information for estimating the centroid position of a single emitter in DD(R). This conclusion can also be extended to DD. With a similar derivation, we have

$$\lim_{d\to\infty} f_{\text{DD},d} = \frac{1}{4} f_{\text{DD},x_0}, \tag{S69}$$

which implies $\sigma_\infty = 2\sigma_{\text{psf}}$, according to the definition in the main text.

#### D. Total Fisher information

Based on the definition of the PSF models in the main text, we can calculate the total information of Polar-SLIVER from all three channels. When $d \to 0$, we have

$$\lim_{d\to 0} f_d = \lim_{d\to 0} f_{\mathrm{R},d} + \frac{1}{2}\lim_{d\to 0} f_{\mathrm{sym},d} + \frac{1}{2}\lim_{d\to 0} f_{\mathrm{asym},d} \tag{S70}$$

$$= \frac{1}{2}\lim_{d\to 0} f_{\mathrm{sym},d} \tag{S71}$$

$$= \sum_{j=\mathrm{x,y}} \left(\frac{\partial \tilde{\omega}_{j\mathrm{A}}}{\partial x}\right)^2. \tag{S72}$$

The above equation shows that the total information at $d \to 0$ entirely comes from the symmetric channel of SLIVER(A). The total photon count of the three channels can be calculated at $d \to 0$ and is

$$N = \sum_{j=\mathrm{x,y,z}} \tilde{\omega}_{j\mathrm{R}}^2 + \sum_{j=\mathrm{x,y}} \tilde{\omega}_{j\mathrm{A}}^2. \tag{S73}$$

In our simulations, $\tilde{\omega}_{j\mathrm{A}}$ and $\tilde{\omega}_{j\mathrm{R}}$ are normalized such that $N = 1$. Considering only x-oriented dipoles, where $j = \mathrm{x}$, we have

$$\lim_{d\to 0} f_d = \left(\frac{\partial \tilde{\omega}_{\mathrm{xA}}}{\partial x}\right)^2. \tag{S74}$$

When $d \to \infty$, we have

$$\lim_{d\to \infty} f_d = \lim_{d\to \infty} f_{\mathrm{R},d} + \frac{1}{2}\lim_{d\to \infty} f_{\mathrm{sym},d} + \frac{1}{2}\lim_{d\to \infty} f_{\mathrm{asym},d} \tag{S75}$$

$$= \lim_{d\to \infty} f_{\mathrm{R},d} + \lim_{d\to \infty} f_{\mathrm{sym},d} \tag{S76}$$

$$= \frac{\left(\sum_{j=\mathrm{x,y,z}} \tilde{\omega}_{j\mathrm{R}} \frac{\partial \tilde{\omega}_{j\mathrm{R}}}{\partial x}\right)^2}{\sum_{j=\mathrm{x,y,z}} \tilde{\omega}_{j\mathrm{R}}^2} + \frac{\left(\sum_{j=\mathrm{x,y}} \tilde{\omega}_{j\mathrm{A}} \frac{\partial \tilde{\omega}_{j\mathrm{A}}}{\partial x}\right)^2}{\sum_{j=\mathrm{x,y}} \tilde{\omega}_{j\mathrm{A}}^2}. \tag{S77}$$

The above equation can be simplified by defining,

$$p_{j\mathrm{R}} = \frac{\tilde{\omega}_{j\mathrm{R}}}{\sqrt{\sum_{j=\mathrm{x,y,z}} \tilde{\omega}_{j\mathrm{R}}^2}}, \quad p_{j\mathrm{A}} = \frac{\tilde{\omega}_{j\mathrm{A}}}{\sqrt{\sum_{j=\mathrm{x,y}} \tilde{\omega}_{j\mathrm{A}}^2}}, \tag{S78}$$

and we have

$$\lim_{d\to \infty} f_d = \left(\sum_{j=\mathrm{x,y,z}} p_{j\mathrm{R}} \frac{\partial \tilde{\omega}_{j\mathrm{R}}}{\partial x}\right)^2 + \left(\sum_{j=\mathrm{x,y}} p_{j\mathrm{A}} \frac{\partial \tilde{\omega}_{j\mathrm{A}}}{\partial x}\right)^2. \tag{S79}$$

From the above equation, we recognize that the total information for estimating $d$ at $d \to \infty$ is exactly one-fourth of the information for estimating the centroid position of a single emitter in DD(A)+DD(R). Considering only one dipole orientation along the x-axis, we have $p_{j\mathrm{R}} = p_{j\mathrm{A}} = 1$ and

$$\lim_{d\to \infty} f_d = \left(\frac{\partial \tilde{\omega}_{\mathrm{xR}}}{\partial x}\right)^2 + \left(\frac{\partial \tilde{\omega}_{\mathrm{xA}}}{\partial x}\right)^2. \tag{S80}$$

Comparing Eq. (S74) with Eq. (S80), the information from SLIVER(A) at $d \to 0$ and $d \to \infty$ are equal, which is consistent with previous results derived from a Gaussian PSF model [14]. However, when considering all dipole orientations, such as for a freely rotating dipole, the information from SLIVER(A) at $d \to 0$ is higher than the information from SLIVER(A) at $d \to \infty$.

#### E. Extension to complex functions

The above derivations assume $\tilde{\omega}_{j\mathrm{A}}$ and $\tilde{\omega}_{j\mathrm{R}}$ are real functions. Under the condition of refractive index matching, $\tilde{\omega}_{j\mathrm{A}}$ is real, but $\tilde{\omega}_{j\mathrm{R}}$ is complex. And when refractive indices are mismatched, both $\tilde{\omega}_{j\mathrm{A}}$ and $\tilde{\omega}_{j\mathrm{R}}$ are complex functions. Take the symmetric channel in SLIVER(A) as an example, we can rewrite Eq. (S40) as

$$I_{\text{2em,sym,A}} = \sum_{j=\text{x,y}} \frac{1}{2} \left| \tilde{\omega}_{j\text{A1}} - \tilde{\omega}_{j\text{A2}} \right|^2 \tag{S81}$$

$$= \sum_{j=\text{x,y}} \frac{1}{2} \left| \tilde{\omega}_{j\text{aA1}} + i\tilde{\omega}_{j\text{bA1}} - \tilde{\omega}_{j\text{aA2}} - i\tilde{\omega}_{j\text{bA2}} \right|^2 \tag{S82}$$

$$= \sum_{j=\text{x,y}} \frac{1}{2} \left( \tilde{\omega}_{j\text{aA1}} - \tilde{\omega}_{j\text{aA2}} \right)^2 + \frac{1}{2} \left( \tilde{\omega}_{j\text{bA1}} - \tilde{\omega}_{j\text{bA2}} \right)^2 \tag{S83}$$

$$= \sum_{\substack{j=\text{x,y} \\ k=\text{a,b}}} \frac{1}{2} \left( \tilde{\omega}_{jk\text{A1}} - \tilde{\omega}_{jk\text{A2}} \right)^2, \tag{S84}$$

where the subscript $k = \text{a}, \text{b}$ denotes the real and imaginary parts of $\tilde{\omega}_{j\text{A}}$. We notice that Eq. (S84) has a similar form to Eq. (S41), with an additional summation over $k$. As previously shown, the summation does not affect the subsequent derivation. Analogous to Eqs. (S47, S53), we can directly write the results at $d \to 0$ and $d \to \infty$ as

$$\lim_{d\to 0} f_{\text{sym},d} = \sum_{\substack{j=\text{x,y} \\ k=\text{a,b}}} 2 \left( \frac{\partial \tilde{\omega}_{\text{jkA}}}{\partial x} \right)^2, \tag{S85}$$

$$\lim_{d\to \infty} f_{\text{sym},d} = \frac{\left( \sum_{\substack{j=\text{x,y} \\ k=\text{a,b}}} \tilde{\omega}_{\text{jkA}} \frac{\partial \tilde{\omega}_{\text{jkA}}}{\partial x} \right)^2}{\sum_{\substack{j=\text{x,y} \\ k=\text{a,b}}} \tilde{\omega}_{\text{jkA}}^2}. \tag{S86}$$

The corresponding results for the antisymmetric channel of SLIVER(A) and for DD(R) can be derived similarly. Below, we directly give the integrands of the total Fisher information for complex functions,

$$\lim_{d\to 0} f_d = \sum_{\substack{j=\text{x,y} \\ k=\text{a,b}}} \left( \frac{\partial \tilde{\omega}_{jk\text{A}}}{\partial x} \right)^2 \tag{S87}$$

$$\lim_{d\to \infty} f_d = \left( \sum_{\substack{j=\text{x,y,z} \\ k=\text{a,b}}} p_{jk\text{R}} \frac{\partial \tilde{\omega}_{jk\text{R}}}{\partial x} \right)^2 + \left( \sum_{\substack{j=\text{x,y} \\ k=\text{a,b}}} p_{jk\text{A}} \frac{\partial \tilde{\omega}_{jk\text{A}}}{\partial x} \right)^2. \tag{S88}$$

where

$$p_{jk\text{R}} = \frac{\tilde{\omega}_{jk\text{R}}}{\sqrt{\sum_{\substack{j=\text{x,y,z} \\ k=\text{a,b}}} \tilde{\omega}_{jk\text{R}}^2}}, \quad p_{jk\text{A}} = \frac{\tilde{\omega}_{jk\text{A}}}{\sqrt{\sum_{\substack{j=\text{x,y} \\ k=\text{a,b}}} \tilde{\omega}_{jk\text{A}}^2}}. \tag{S89}$$

#### F. Extension to pixel integration

The above derivations assume that $\tilde{\omega}_{jk\text{A}}$ and $\tilde{\omega}_{jk\text{R}}$ are continuous functions, and the Fisher information is calculated from a continuous integral of $f_d$. However, in practice, camera detection is pixelated, and the Fisher information is calculated from discrete integration. To incorporate pixel integration, we simulate the pixel value as the sum of the values from four upsampled positions around the pixel coordinates. This integration introduces an additional summation to the two-emitter models. Here, we skip the derivation and directly give the integrands of the total

Fisher information for complex functions under pixelated detection,

$$\lim_{d\to 0} f_{dq} = \sum_{\substack{j=\mathrm{x,y}\\k=\mathrm{a,b}}} \sum_{m} \left( \frac{\partial \tilde{\omega}_{jkqm\mathrm{A}}}{\partial x_{qm}} \right)^2 \tag{S90}$$

$$\lim_{d\to \infty} f_{dq} = \left( \sum_{\substack{j=\mathrm{x,y,z}\\k=\mathrm{a,b}}} \sum_{m} p_{jkqm\mathrm{R}} \frac{\partial \tilde{\omega}_{jkqm\mathrm{R}}}{\partial x_{qm}} \right)^2 + \left( \sum_{\substack{j=\mathrm{x,y}\\k=\mathrm{a,b}}} \sum_{m} p_{jkqm\mathrm{A}} \frac{\partial \tilde{\omega}_{jkqm\mathrm{A}}}{\partial x_{qm}} \right)^2, \tag{S91}$$

$$p_{jkqm\mathrm{R}} = \frac{\tilde{\omega}_{jkqm\mathrm{R}}}{\sqrt{\sum_{\substack{j=\mathrm{x,y,z}\\k=\mathrm{a,b}}} \sum_m \tilde{\omega}^2_{jkqm\mathrm{R}}}}, \tag{S92}$$

$$p_{jkqm\mathrm{A}} = \frac{\tilde{\omega}_{jkqm\mathrm{A}}}{\sqrt{\sum_{\substack{j=\mathrm{x,y}\\k=\mathrm{a,b}}} \sum_m \tilde{\omega}^2_{jkqm\mathrm{A}}}}. \tag{S93}$$

where the subscript $m = 1, 2, 3, 4$ denotes the four upsampled positions around the camera pixel $q$.

### 6. QUANTUM FISHER INFORMATION OF SEPARATION ESTIMATION OF TWO FREELY ROTATING DIPOLES

In this section, we derive the quantum Fisher information (QFI) of estimating the separation between two freely rotating dipole emitters. We assume that the two dipole emitters are centered at the optical axis and are separated by a distance $d$ along the x-axis. The calculation of QFI can be performed either at the pupil plane or the image plane. The results should be equivalent, as the Fourier transform shall not alter the QFI. The signal at the pupil plane is contained in a finite circular region, while the signal at the image plane is extended to infinity. Therefore, we choose to calculate the QFI at the pupil plane. The electric field at the pupil plane from a fixed dipole emitter can be represented by a vector field,

$$\boldsymbol{\psi}(k_x, k_y) = h_{\mathrm{H}}(k_x, k_y)\mathbf{H} + h_{\mathrm{V}}(k_x, k_y)\mathbf{V}, \tag{S94}$$

where $h_{\mathrm{H}}$ and $h_{\mathrm{V}}$ are given by Eqs. (S16, S17), describing the electric field at the pupil plane in horizontal and vertical polarizations, respectively. The intensity field can be calculated from,

$$\boldsymbol{I}(k_x, k_y) = |\boldsymbol{\psi}(k_x, k_y)|^2 = |h_{\mathrm{H}}(k_x, k_y)|^2 + |h_{\mathrm{V}}(k_x, k_y)|^2. \tag{S95}$$

We can also express Eqs. (S94, S95) in terms of vector representation,

$$\boldsymbol{\psi}(k_x, k_y) = \begin{bmatrix} h_{\mathrm{H}}(k_x, k_y) \\ h_{\mathrm{V}}(k_x, k_y) \end{bmatrix}, \tag{S96}$$

$$\boldsymbol{I}(k_x, k_y) = \boldsymbol{\psi}^*(k_x, k_y) \cdot \boldsymbol{\psi}(k_x, k_y), \tag{S97}$$

where the symbol $*$ denotes complex conjugate and the symbol $\cdot$ denotes inner product between two vectors.

Given a set of parameters $\boldsymbol{\theta} = [\theta_1, \theta_2, ..., \theta_N]$, the QFI is defined by [15],

$$\mathcal{K}_{ab} = \frac{1}{2} \mathrm{Tr}\{\rho(L_a L_b + L_b L_a)\}, \tag{S98}$$

where $\rho = \rho(\boldsymbol{\theta})$ is the density operator of the quantum state describing a quantum system, and the parameters are encoded in $\rho$. And $L_a(L_b)$ is the symmetric logarithmic derivative (SLD) for the parameter $\theta_a(\theta_b)$, where $a(b)$ denotes the $a$th ($b$th) parameter. To compute $\mathcal{K}_{ab}$, the standard method is to decompose $\rho$ in terms of its eigenstates,

$$\rho = \sum_i \lambda_i |e_i\rangle \langle e_i|, \tag{S99}$$

where $\lambda_i$ is the corresponding eigenvalue of the eigenstate $|e_i\rangle$. Then QFI can be calculated from [15],

$$\mathcal{K}_{ab} = \sum_{i,j;\lambda_i+\lambda_j\neq 0} \frac{2\mathrm{Re}\left( \langle e_i|\frac{\partial\rho}{\partial\theta_a}|e_j\rangle \langle e_j|\frac{\partial\rho}{\partial\theta_b}|e_i\rangle \right)}{\lambda_i+\lambda_j}. \tag{S100}$$

In the following sections, we will apply the vector fields in the calculation of QFI.

#### A. Density operator of two freely rotating dipoles

In our case, a fluorescent dipole emitter can be considered as a weak thermal source, where the average photon count $\epsilon$ within a coherence time interval is much smaller than one [14]. The density operator of a weak thermal source can be approximated to [14],

$$\rho \approx (1-\epsilon)\,|\mathrm{vac}\rangle\,\langle\mathrm{vac}| + \epsilon\rho_1. \tag{S101}$$

Here, $|\mathrm{vac}\rangle$ denotes the zero-photon state, which does not contribute to the QFI. We consider that the information is mainly from the one-photon state $\rho_1$. For two freely rotating dipoles, the one-photon state can be expressed as[16],

$$\rho_1 = \frac{1}{2}\sum_{s=\mathrm{x,y,z}} |\psi_{1s}\rangle\,\langle\psi_{1s}| + |\psi_{2s}\rangle\,\langle\psi_{2s}|\,, \tag{S102}$$

where 1 and 2 denote emitter 1 and emitter 2. And $|\psi_{1s(2s)}\rangle$ denotes the quantum state of the dipole emitter 1(2) oscillating along the $s$ direction. Now we apply the vector field defined in Eq. (S96) and explain the physical meaning of the density operator. For direct measurement of the intensity field at the pupil plane, the probability of detecting one photon at position $(k_x, k_y)$ is given by

$$\langle (k_x,k_y)|\rho_1|(k_x,k_y)\rangle = \frac{1}{2}\sum_{s=\mathrm{x,y,z}} \boldsymbol{I}_{1s}(k_x,k_y) + \boldsymbol{I}_{2s}(k_x,k_y), \tag{S103}$$

where $|(k_x,k_y)\rangle$ is the eigenket for a pupil-plane position, and $\boldsymbol{I}_{1s(2s)}(k_x,k_y)$ can be calculated from Eq. (S97). The state $|\psi_{1s(2s)}\rangle$ can be expressed as an expansion of $|(k_x,k_y)\rangle$,

$$|\psi_{1s(2s)}\rangle = \iint dk_x dk_y \boldsymbol{\psi}_{1s(2s)}(k_x,k_y)\,|(k_x,k_y)\rangle\,. \tag{S104}$$

The inner product of the two pure states $\langle\psi_u|\psi_v\rangle$ is given by

$$\langle\psi_u|\psi_v\rangle = \iint dk_x dk_y \boldsymbol{\psi}_u^*(k_x,k_y)\cdot\boldsymbol{\psi}_v(k_x,k_y), \quad u,v = 1s, 2s, \quad s = \mathrm{x,y,z}. \tag{S105}$$

The state $|\psi_s\rangle$ is normalized so that

$$\sum_{s=\mathrm{x,y,z}} \langle\psi_s|\psi_s\rangle = 1. \tag{S106}$$

The vector fields of emitter 1 and emitter 2, separated by a distance $d$, are defined as follows:

$$\boldsymbol{\psi}_{1s}(k_x,k_y,d) = \boldsymbol{\psi}_s(k_x,k_y)\exp\left(-ik_x\frac{d}{2}\right), \tag{S107}$$

$$\boldsymbol{\psi}_{2s}(k_x,k_y,d) = \boldsymbol{\psi}_s(k_x,k_y)\exp\left(ik_x\frac{d}{2}\right), \tag{S108}$$

#### B. Eigen decomposition of $\rho_1$

To compute the QFI from Eq. (S100), we need to find the eigenstates of $\rho_1$, such that

$$\rho_1 = \sum_i \lambda_i\,|e_i\rangle\,\langle e_i|\,, \tag{S109}$$

Finding the analytical expression of $|e_i\rangle$ can be complicated. Here, we numerically calculate $|e_i\rangle$ at a given separation $d$. In numerical calculation, we consider $|\psi_s\rangle$ as a 2D array, generated from evenly spaced samples of $\boldsymbol{\psi}_s(k_x,k_y)$ in the pupil plane. From Eq. (S96), each element of $|\psi_s\rangle$ is a vector,

$$\boldsymbol{\psi}_{sp} = \begin{bmatrix} h_{\mathrm{H},s}(k_{xp},k_{yp}) \\ h_{\mathrm{V},s}(k_{xp},k_{yp}) \end{bmatrix}, \quad s = \mathrm{x,y,z} \tag{S110}$$

where $p$ indexes the 2D array. We then compute the inner product of two pure states as,

$$\langle \psi_u | \psi_v \rangle = \sum_p \boldsymbol{\psi}_{up}^* \cdot \boldsymbol{\psi}_{vp}, \quad u, v = 1s, 2s, \quad s = \mathrm{x, y, z}. \tag{S111}$$

To find the eigenstates, we first compute the inner product matrix of a set of six pure states, $\boldsymbol{\Psi} = [|\psi_{1x}\rangle, |\psi_{1y}\rangle, |\psi_{1z}\rangle, |\psi_{2x}\rangle, |\psi_{2y}\rangle, |\psi_{2z}\rangle]^T$ as,

$$M_{uv} = \langle \psi_u | \psi_v \rangle, \quad u, v = 1s, 2s, \quad s = \mathrm{x, y, z}. \tag{S112}$$

Here, $T$ denotes transpose and $M$ is a $6 \times 6$ complex matrix. We then apply singular value decomposition (SVD) to $M$,

$$M = USV^\dagger, \tag{S113}$$

where the symbol $\dagger$ denotes conjugate transpose. The decomposition of $\boldsymbol{\Psi}$ in terms of eigen basis can be expressed as,

$$\boldsymbol{\Psi} = A\boldsymbol{e}, \tag{S114}$$

where $A$ is a $6 \times 6$ complex matrix and $\boldsymbol{e} = [|e_1\rangle, |e_2\rangle, |e_3\rangle, |e_4\rangle, |e_5\rangle, |e_6\rangle]^T$ is a vector of six eigenstates. We can obtain $A$ from

$$A = \left(\sqrt{S}V^\dagger\right)^T. \tag{S115}$$

Then the eigenstates can be constructed from

$$\boldsymbol{e} = A^{-1}\boldsymbol{\Psi}. \tag{S116}$$

Substituting Eq. (S114) into Eq. (S102), we can obtain the coefficient $C_{ij}$ of each term $|e_i\rangle \langle e_j|$ from,

$$C_{ij} = \frac{1}{2}(A^\dagger A)_{ij}. \tag{S117}$$

According to Eq. (S115), the matrix $A^\dagger A$ can be further calculated as,

$$A^\dagger A = \sqrt{S}V^T V^* \sqrt{S} = S. \tag{S118}$$

Since the matrix $S$ is a real diagonal matrix, we have $C_{ij} = 0, i \neq j$. With only diagonal terms, we obtain the eigen decomposition of $\rho_1$ as

$$\rho_1 = \sum_i C_{ii} |e_i\rangle \langle e_i|. \tag{S119}$$

Let $\lambda_i = C_{ii}$, the eigenvalues are,

$$\boldsymbol{\lambda} = \frac{1}{2}\mathrm{diag}\,(S), \tag{S120}$$

where diag() denotes the operator of taking the diagonal elements from a matrix, and $\boldsymbol{\lambda} = [\lambda_1, \lambda_2, \lambda_3, \lambda_4, \lambda_5, \lambda_6]^T$ is a vector of six eigenvalues. Now we find six eigenstates that can fully describe $\rho_1$.

From Eq. (S100), we need to calculate the derivative of $\rho_1$ with respect to the estimation parameter. In our case of a single estimation parameter, separation $d$, we have

$$\frac{\mathrm{d}\rho_1}{\mathrm{d}d} = \frac{1}{2}\sum_s \frac{\mathrm{d}\,|\psi_{1s}\rangle}{\mathrm{d}d}\langle \psi_{1s}| + |\psi_{1s}\rangle \frac{\mathrm{d}\,\langle \psi_{1s}|}{\mathrm{d}d} + \frac{\mathrm{d}\,|\psi_{2s}\rangle}{\mathrm{d}d}\langle \psi_{2s}| + |\psi_{2s}\rangle \frac{\mathrm{d}\,\langle \psi_{2s}|}{\mathrm{d}d}. \tag{S121}$$

Let

$$|\psi'_{1s}\rangle = \frac{\mathrm{d}\,|\psi_{1s}\rangle}{\mathrm{d}d}, \quad |\psi'_{2s}\rangle = \frac{\mathrm{d}\,|\psi_{2s}\rangle}{\mathrm{d}d}. \tag{S122}$$

According to Eqs. S107,S108), we have

$$\boldsymbol{\psi}'_{1s}(k_x, k_y) = \frac{\mathrm{d}\boldsymbol{\psi}_{1s}}{\mathrm{d}d} = -\frac{ik_x}{2}\boldsymbol{\psi}_{1s}, \tag{S123}$$

$$\boldsymbol{\psi}'_{2s}(k_x, k_y) = \frac{\mathrm{d}\boldsymbol{\psi}_{2s}}{\mathrm{d}d} = \frac{ik_x}{2}\boldsymbol{\psi}_{2s}. \tag{S124}$$

We can again construct $|\psi'_{1s}\rangle$ and $|\psi'_{2s}\rangle$ as 2D arrays from evenly spaced samples of $\boldsymbol{\psi}'_{1s}(k_x, k_y)$ and $\boldsymbol{\psi}'_{2s}(k_x, k_y)$, respectively. It can be shown that $|\psi'_{1s}\rangle$ and $|\psi'_{2s}\rangle$ cannot be fully expanded by the six found eigenstates. We need additional dimensions to support the space spanning $|\psi'_{1s}\rangle$

and $|\psi'_{2s}\rangle$ [14]. To achieve this, we first remove from $|\psi'_{1s(2s)}\rangle$ the projection of itself in the space of $\boldsymbol{e}$ as follows,

$$p_{ui} = \langle e_i | \psi'_u \rangle, \quad i = 1, 2, ..., 6, \quad u = 1s, 2s, \quad s = \mathrm{x, y, z}, \tag{S125}$$

$$|\psi'_{u\perp}\rangle = |\psi'_u\rangle - \sum_i p_{ui} |e_i\rangle, \tag{S126}$$

where each inner product is calculated according to Eq. (S111). The above operation is repeated until the sum $\sum_i p^2_{ui}$ for all $u$ is less than a set tolerance. Then we follow the same procedure as in Eqs. (S112-S116) to find another set of six eigenstates $\boldsymbol{e}' = [|e_7\rangle, |e_8\rangle, |e_9\rangle, |e_{10}\rangle, |e_{11}\rangle, |e_{12}\rangle]^T$, by replacing $\boldsymbol{\Psi}$ with $\boldsymbol{\Psi}'$, where

$$\boldsymbol{\Psi}' = \left[ |\psi'_{1x\perp}\rangle, |\psi'_{1y\perp}\rangle, |\psi'_{1z\perp}\rangle, |\psi'_{2x\perp}\rangle, |\psi'_{2y\perp}\rangle, |\psi'_{2z\perp}\rangle \right]^T. \tag{S127}$$

The eigenvalues for $\boldsymbol{e}'$ are all zero, which is $\lambda_i = 0, i = 7, \ldots, 12$. We now find 12 eigenstates and their corresponding eigenvalues.

#### C. Calculation of QFI

Given the eigen decomposition of $\rho_1$, substituting Eq. (S121) into Eq. (S100), we can compute the QFI of estimating $d$ from,

$$\mathcal{K}_d = \sum_{\substack{i,j=1 \\ \lambda_i + \lambda_j \neq 0}}^{12} \frac{2\mathrm{Re}\left( \langle e_i | \frac{\mathrm{d}\rho_1}{\mathrm{d}d} | e_j \rangle \langle e_j | \frac{\mathrm{d}\rho_1}{\mathrm{d}d} | e_i \rangle \right)}{\lambda_i + \lambda_j}. \tag{S128}$$

where each inner product is calculated according to Eq. (S111). To illustrate the above derivation, we next give an example of calculating $\mathcal{K}_d$. Assuming that two freely rotating dipole emitters are separated by $d = 0.01\sigma_{\mathrm{psf}}$, based on Eq. (S110), we can obtain six vector fields at the pupil plane from the two dipole emitters. Figure S11(a) shows the three vector fields from emitter 1. To visualize the vector field, we rewrite Eq. (S96) as,

$$\boldsymbol{\psi} = \begin{bmatrix} |h_{\mathrm{H}}| \\ |h_{\mathrm{V}}| e^{i\Delta\varphi} \end{bmatrix} e^{i\varphi_{\mathrm{H}}}. \tag{S129}$$

Here, we omit $(k_x, k_y)$ for simplicity. The phase delay between horizontal and vertical polarizations is $\Delta\varphi = \varphi_{\mathrm{V}} - \varphi_{\mathrm{H}}$. The field inside the bracket defines the polarization state at $(k_x, k_y)$: when $\Delta\varphi = 0$ or $\pi$, the field is linearly polarized; otherwise, the field is elliptically polarized. We can define the ellipse as

$$k_x = a|h_{\mathrm{H}}| \cos t, \tag{S130}$$

$$k_y = a|h_{\mathrm{V}}| \cos(t + \Delta\varphi), \tag{S131}$$

where $a$ is a scaling factor to adjust the size of the ellipse for visualization, and $t$ is evenly sampled from 0 to $2\pi$. For an elliptically polarized field, when $\pi < \Delta\varphi < 2\pi$, the polarization is right-handed, and when $0 < \Delta\varphi < \pi$, it is left-handed.

Following the eigen decomposition of $\rho_1$ in section 6B, we obtain twelve eigenstates at the given separation. To quantify the contribution of each eigenstate to $\mathcal{K}_d$, we rewrite Eq. (S128) as

$$\mathcal{K}_d = \sum_{i,j=1}^{12} L_{ij}. \tag{S132}$$

Figure S11(c) shows the resulting $12 \times 12$ matrix constructed from $L_{ij}$. The elements with the highest values are $L_{44}, L_{55}, L_{66}$, which are contributed by eigenstates $|e_4\rangle, |e_5\rangle, |e_6\rangle$ [Fig. S11(a)]. We found that at $d < 0.5\sigma_{\mathrm{psf}}$, most information is contributed by those three eigenstates [Fig. S11(c,d)]. According to Eq. (S116), we can explicitly calculate those eigenstates at small separations as,

$$|e_4\rangle = a_{4\mathrm{x}} \left( |\psi_{1\mathrm{x}}\rangle + |\psi_{2\mathrm{x}}\rangle \right) + a_{4\mathrm{z}} \left( |\psi_{1\mathrm{z}}\rangle - |\psi_{2\mathrm{z}}\rangle \right), \tag{S133}$$

$$|e_5\rangle = a_{5\mathrm{y}} \left( |\psi_{1\mathrm{y}}\rangle - |\psi_{2\mathrm{y}}\rangle \right), \tag{S134}$$

$$|e_6\rangle = a_{6\mathrm{x}} \left( |\psi_{1\mathrm{x}}\rangle - |\psi_{2\mathrm{x}}\rangle \right) + a_{6\mathrm{z}} \left( |\psi_{1\mathrm{z}}\rangle + |\psi_{2\mathrm{z}}\rangle \right). \tag{S135}$$

The coefficients $a_{ij}, i = 4, 5, 6, j = \mathrm{x}, \mathrm{y}, \mathrm{z}$ are complex numbers and are dependent on separation. We found that the relative magnitudes of those coefficients remain nearly unchanged with respect to $d$ when $d < 0.5\sigma_{\mathrm{psf}}$ [Fig. S11(b)]. The dominant coefficients are $a_{6\mathrm{x}}, a_{5\mathrm{y}}, a_{4\mathrm{z}}$, corresponding to the terms $|\psi_{1s}\rangle - |\psi_{2s}\rangle, s = \mathrm{x}, \mathrm{y}, \mathrm{z}$, respectively. In quantum information theory, the optimal measurement is given by the eigenstates when the eigenstates are independent of the estimation parameters [15]. In our case, the estimation parameter is $d$. At small separations, we observe that the dominant eigenstates are nearly independent of $d$ [Fig. S11(b)]. This suggests that the optimal measurement in this regime should measure the intensities from $|\psi_{1s}\rangle - |\psi_{2s}\rangle, s = \mathrm{x}, \mathrm{y}, \mathrm{z}$. However, implementing such a measurement in practice is challenging.

As we demonstrated in the main text, the electric fields of x- and y-oriented dipoles exhibit opposite symmetry compared to that of a z-oriented dipole. When projecting the fields onto the azimuthal polarization, the contribution from the z-oriented dipole vanishes. Therefore, the SLIVER(A) channel can access partial information from $|\psi_{1s}\rangle - |\psi_{2s}\rangle, s = \mathrm{x}, \mathrm{y}$. However, the radial polarization contains contributions from x-, y-, and z-oriented dipoles, making it impossible to measure the intensity from $|\psi_{1\mathrm{z}}\rangle - |\psi_{2\mathrm{z}}\rangle$. Therefore, our proposed measurement, Polar-SLIVER, is suboptimal for measuring the separation of two freely rotating dipoles. Achieving the quantum-optimal measurement would require techniques capable of isolating the z-oriented dipole contribution from those of the x- and y-oriented dipoles.

#### D. Comparison of QFIs between dipole emitters and scalar point sources

For a scalar point source, the electric field at the pupil plane is considered as a scalar field without polarization. For example, a linearly polarized field can be considered as a scalar field. For a microscope system, the simplest pupil can be described by a circle function [17],

$$h = \mathrm{circ}\left(\frac{\sqrt{k_x^2 + k_y^2}}{n_a/\lambda}\right), \tag{S136}$$

where $n_a/\lambda$ defines the microscope's cut-off spatial frequency, with $n_a$ being the numerical aperture of the objective and $\lambda$ the emission wavelength. The PSF of such pupil function is the Airy pattern. Shifting the scalar point source along the x-axis adds a phase term to the pupil function,

$$h(x) = \mathrm{circ}\left(\frac{\sqrt{k_x^2 + k_y^2}}{n_a/\lambda}\right) \exp(ik_x x). \tag{S137}$$

The partial derivative of $h(x)$ with respect to $x$ is

$$\frac{\partial h(x)}{\partial x} = h(x) 2\pi i k_x. \tag{S138}$$

The QFI for estimating the separation of two scalar point sources is given by [14],

$$\mathcal{K}_{d,\mathrm{scalar}} = \frac{\iint \left|\frac{\partial h(x)}{\partial x}\right|^2 dk_x dk_y}{\iint |h(x)|^2 dk_x dk_y}. \tag{S139}$$

The normalization term ensures that the photon count is one. Substituting Eqs. (S137, S138) into Eq. (S139), we have

$$\mathcal{K}_{d,\mathrm{Airy}} = \frac{\pi^2 n_a^2}{\lambda^2}. \tag{S140}$$

The above result is consistent with the result obtained by approximating the Airy pattern as a Gaussian PSF and letting $\sigma_{\mathrm{psf}} = \lambda/(2\pi n_a)$ [14, 18].

We compared the QFIs for estimating the separation of two dipole emitters and two Airy patterns (Fig. S12). For a freely rotating dipole, the electric field includes contributions from x-, y- and z-oriented dipoles [Fig. S12(a,c)]. We found that as $d \to \infty$, $\mathcal{K}_{d,\mathrm{dipole}}$ approaches $\mathcal{K}_{d,\mathrm{Airy}}$ when NA<0.2. However, as $d \to 0$, $\mathcal{K}_{d,\mathrm{dipole}}$ continuously decreases below $\mathcal{K}_{d,\mathrm{Airy}}$ with decreasing NA, and only at extremely low NA (e.g., 0.001) does it converge back to $\mathcal{K}_{d,\mathrm{Airy}}$.

When the z-oriented dipole is removed, we found that $\mathcal{K}_{d,\mathrm{dipole}}$ becomes independent of separation and is always above $\mathcal{K}_{d,\mathrm{Airy}}$ [Fig. S12(b,d)]. Furthermore, $\mathcal{K}_{d,\mathrm{dipole}}$ approaches $\mathcal{K}_{d,\mathrm{Airy}}$ at NA<0.9. This is because at low NAs, the electric fields at the pupil plane from x- and y-oriented dipoles can be approximated to a linearly polarized field. Since their fields are nearly orthogonal, the QFI can be approximated by the sum of the QFIs from independent scalar fields. Therefore, at low NAs, the QFI of x- and y-oriented dipoles approaches that of Airy patterns.

However, this approximation is only valid at ultra-low NAs for freely rotating dipoles. The z-oriented dipole produces a radially polarized field at the pupil plane, which cannot be approximated as linearly polarized at any NA. Therefore, when all three dipole components (x, y, and z) are present, we cannot approximate the electric field at the pupil plane to a scalar field, except for ultra-low NAs, where the z-oriented dipole contribution is negligible.
